# Sequence-based machine learning reveals 3D genome differences between bonobos and chimpanzees

**DOI:** 10.1101/2023.10.26.564272

**Authors:** Colin M. Brand, Shuzhen Kuang, Erin N. Gilbertson, Evonne McArthur, Katherine S. Pollard, Timothy H. Webster, John A. Capra

## Abstract

Phenotypic divergence between closely related species, including bonobos and chimpanzees (genus *Pan*), is largely driven by variation in gene regulation. The 3D structure of the genome mediates gene expression; however, genome folding differences in *Pan* are not well understood. Here, we apply machine learning to predict genome-wide 3D genome contact maps from DNA sequence for 56 bonobos and chimpanzees, encompassing all five extant lineages. We use a pairwise approach to estimate 3D divergence between individuals from the resulting contact maps in 4,420 1 Mb genomic windows. While most pairs were similar, *∼*17% were predicted to be substantially divergent in genome folding. The most dissimilar maps were largely driven by single individuals with rare variants that produce unique 3D genome folding in a region. We also identified 89 genomic windows where bonobo and chimpanzee contact maps substantially diverged, including several windows harboring genes associated with traits implicated in *Pan* phenotypic divergence. We used *in silico* mutagenesis to identify 51 3D-modifying variants in these bonobo-chimpanzee divergent windows, finding that 34 or 66.67% induce genome folding changes via CTCF binding motif disruption. Our results reveal 3D genome variation at the population-level and identify genomic regions where changes in 3D folding may contribute to phenotypic differences in our closest living relatives.

## 1 Introduction

Phenotypic divergence between closely related species is largely driven by variation in gene regulation, including humans and our closest living relatives (Enard et al., 2002; King and Wilson, 1975; Sholtis and Noonan, 2010). The three-dimensional (3D) organization of the genome is increasingly recognized as a key mediator of gene expression by facilitating interactions between distal and proximal cis-regulatory elements (Bonev and Cavalli, 2016; Dekker et al., 2023; Ibrahim and Mundlos, 2020). Consequently, disruption of genome folding has been associated with human disease (Lupiáñez et al., 2015; Norton and Phillips-Cremins, 2017) and variation in genome folding underlies traits that differ between humans and other species (Batyrev et al., 2020; Keough et al., 2022; McArthur et al., 2022).

Humans’ closest living relatives, bonobos (*Pan paniscus*) and chimpanzees (*P. troglodytes*), exhibit a number of phenotypic differences (Stumpf, 2011; Gruber and Clay, 2016); yet, the molecular mechanisms that contribute to this divergence remain elusive. Species-specific protein differences identified from missense single nucleotide variants (SNVs) in population-level genomic data (de Manuel et al., 2016; Prado-Martinez et al., 2013) are the most well characterized (Cagan et al., 2016; Han et al., 2019; Kovalaskas et al., 2020; Prüfer et al., 2012). In contrast, gene regulatory differences between bonobos and chimpanzees are less understood and primarily studied in the context of human uniqueness (Khrameeva et al., 2020; Marchetto et al., 2013). Understanding gene regulation in *Pan* is further impeded by limited -omics data, especially data from assays of 3D genome folding such as Hi-C and Micro-C (Kempfer and Pombo, 2020). Currently, there are publicly available Hi-C samples from four chimpanzee induced pluripotent stem cells, one chimpanzee lymphoblastoid cell line (LCL), and one bonobo LCL (Eres et al., 2019; Yang et al., 2019).

Here, we leverage a machine learning algorithm that predicts 3D genome folding from DNA sequence (Fudenberg et al., 2020) to assess the contribution of the 3D genome to regulatory variation in bonobos and chimpanzees at population-scale. First, we evaluate model performance on chimpanzee sequence and describe the generation of chromatin contact maps. Second, we assess inter-individual variation in chromatin contact genome-wide and among smaller genomic windows. Third, we identify windows that exhibit species-specific genome folding, some of which harbor genes with species differences in gene expression. Fourth, we discover individual variants that drive genome folding differences between species. These results provide a foundation for exploring this essential gene regulatory mechanism in our closest living relatives.

## 2 Results

### 2.1 Akita predicts genome folding in bonobos and chimpanzees

We first characterized the performance of Akita (Fudenberg et al., 2020), a deep learning algorithm that predicts chromatin contact from DNA sequence, on a chimpanzee genome. Akita predicts 3D contacts from 1,048,576 bp of sequence, estimating contacts for the center 917,504 bp of a given window at 2,048 bp resolution. Akita was simultaneously trained on Hi-C and Micro-C datasets from humans and performed reasonably well when applied to mice (median Spearman *ρ* = 0.50) (Fudenberg et al., 2020). We used chimpanzee sequence to generate predictions for the human foreskin fibroblast (HFF) cell type and compared to chimpanzee neural progenitor cell (NPC) Hi-C data (**Figure S1A**). The predictions accurately capture the main structural patterns of chimpanzee 3D genome (held-out test set regions: Spearman *ρ ∼* 0.44) (**Figures S1B**, **S1C**). The model has lowest accuracy on regions of the chimpanzee genome with minimally consistent 3D structure—regions that have low correlations in human data.

We thus examined differences in 3D organization among *Pan* lineages by predicting genome-wide 3D contact maps for individuals from all five extant lineages (**Figure 1A**). We identified high-quality genotypes for single nucleotide variants (SNVs) called (Brand et al., 2021) from data generated for 71 individuals (de Manuel et al., 2016; Prado-Martinez et al., 2013). This procedure resulted in 1,137,208 to 9,393,495 SNVs per individual (**File S1**, **Figure S2**). Next, we inserted each individual’s set of SNVs into the chimpanzee reference sequence, panTro6 (Kronenberg et al., 2018) (**File S1**). After filtering individuals with low-quality genotypes, we retained 56 individuals for downstream analyses: nine bonobos, five Nigeria-Cameroon chimpanzees, 17 eastern chimpanzees, 16 central chimpanzees, and nine western chimpanzees (**Figures 1A**, **S2**, **File S1**). We tiled the chimpanzee reference genome with 5,317 sliding windows that overlapped by half. We discarded windows without complete sequence coverage (i.e., *≥* 1 “N”s), retaining 4,420 windows. We applied Akita to sequences for all 56 individuals at these full-coverage windows (**Figure 1B**).

**Figure 1:**
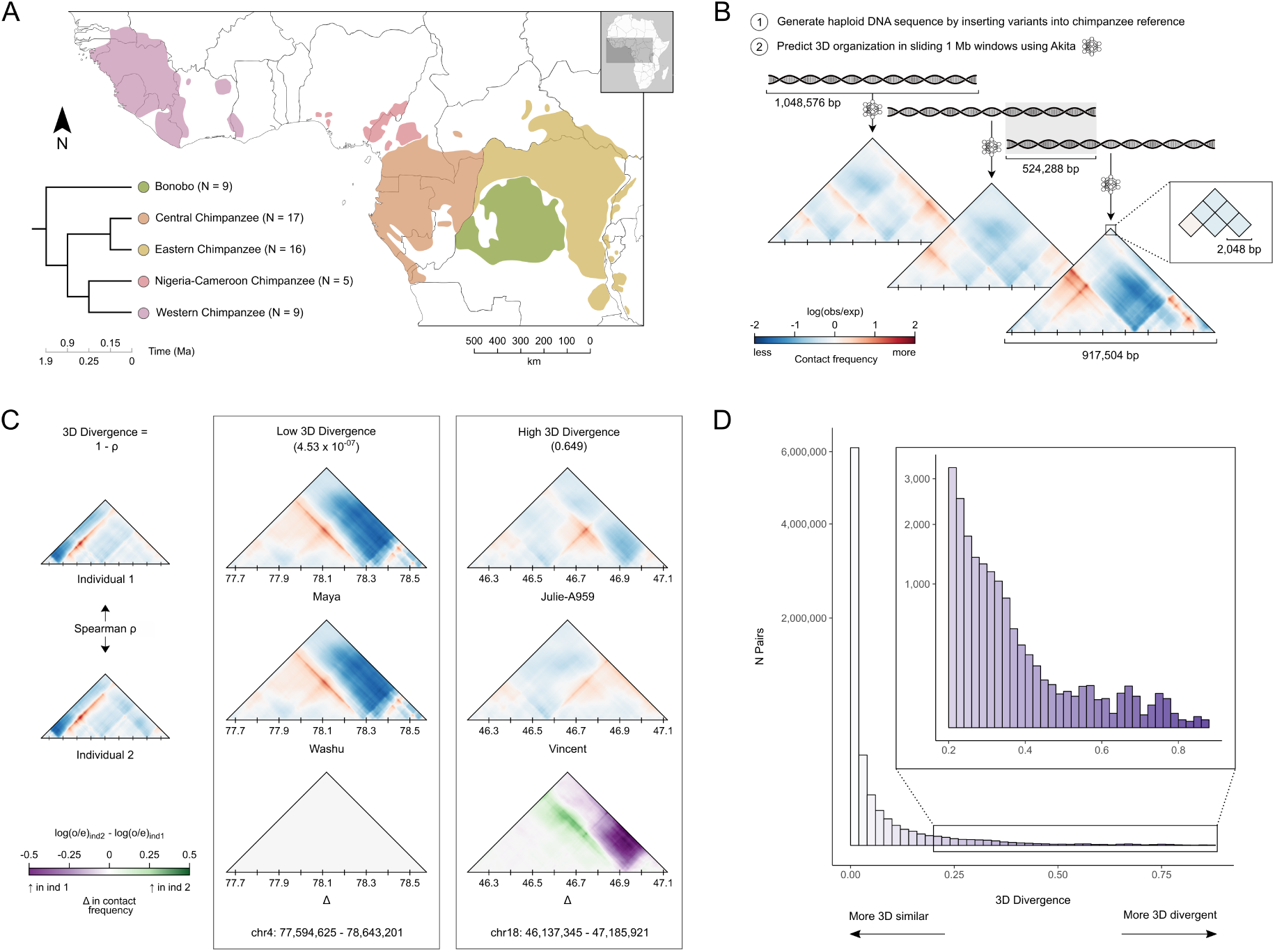
Predicting genome folding from DNA sequence in bonobos and chimpanzees. **(A)** The geographic distribution and evolutionary relationships among all five extant *Pan* lineages. Divergence times are from Brand et al., 2022; de Manuel et al., 2016. Ns indicate the sample size after filtering individuals with lowquality genotypes. **(B)** Schematic of the generation of genome-wide 3D contact maps for 56 *Pan* individuals. We inserted the single nucleotide variants from each individual into the chimpanzee reference DNA sequence (panTro6) and then applied Akita to each sequence. Akita takes 1,048,576 bp of DNA sequence as input and generates a 3D contact map for the central 917,504 bp of the window. The map consists of predicted contacts for all pairs of 2,048 bp loci within the window. We applied Akita to sliding windows overlapping by half across the genome resulting in 5,317 windows. We discarded windows without full sequence coverage in the reference sequence, yielding 4,420 analyzable windows. **(C)** Example comparisons of 3D genome divergence in the contact maps between pairs of individuals. To quantify divergence, we calculated “3D divergence” as the Spearman correlation coefficient over the corresponding cells for a given pair of maps subtracted from 1. Thus, as illustrated, a divergence score near 0 indicates high similarity, whereas greater divergence scores indicate dissimilarity. Contact frequencies per cell for the individual maps are colored as in **B**. The Δ map illustrates the contact frequency difference for the pair (individual 2 - individual 1). **(D)** The distribution of 3D divergence. We compared all pairs of individuals for all autosomal windows, resulting in a total of 6,541,920 pairs. We also compared contact maps for the X chromosome among all pairs of females (N = 95,130). Scores are binned using 0.02 steps from 0 to 0.88. The inset shows divergence > 0.2. Note the y-axis is square root transformed.

To quantify divergence in predicted contact maps genome-wide, we compared all autosomal windows between all pairs of individuals (N = 6,541,920) (**Figure 1C**). We restricted our analysis of X chromosome windows to pairs of females (N = 95,130) because the chromosome is hemizygous in males. We calculated “3D divergence” as 1 - *ρ* for all pixels per pair of maps (**Figure 1C**) (McArthur et al., 2022). Lower 3D divergence indicates greater similarity in contact maps, whereas higher 3D divergence suggests contact map differences (**Figure 1C**). We use 1 - *ρ* here because this map comparison method is sensitive to map differences due to structural differences yet agnostic to differences in contact frequency (Gunsalus et al., 2023b), enabling us to focus on 3D structural differences in *Pan* in this analysis. Hereafter, “window” indicates any of the 4,420 1 Mb windows used in the analysis and “pair” denotes a comparison between two contact maps of a given window for two different individuals.

### 2.2 3D genome folding is largely conserved across bonobos and chimpanzees

We first summarized patterns of 3D contact map similarity and divergence across all pairs of individuals and all genomic windows. Based on previous work indicating substantial evolutionary constraint on genome folding (Fudenberg and Pollard, 2019; Krefting et al., 2018; McArthur and Capra, 2021), we anticipated that most pairs would exhibit minimal divergence. As expected, most contact maps were extremely similar between pairs of individuals (**Figure 1D**), including 5,539,567 pairs or 83.06% which had 3D divergence < 0.01.

To explore this conservation in a deeper phylogenetic context with experimental data, we analyzed conserved topologically associated domains (TADs) identified from Hi-C data generated from four murine and four primate species (Okhovat et al., 2023). We quantified patterns of 3D divergence among *Pan* individuals in these deeply conserved regions. Windows intersecting TADs conserved across the four primate species had significantly lower 3D divergence (mean maximum of 0.0332) than the divergence observed genome-wide (mean maximum of 0.0502; Komologorov-Smirnov, K = 0.13, P = 8.71 *×* 10^-6^) (**Figure S3A**). The divergence was even smaller for pairs intersecting “ultraconserved” TAD boundaries observed in all eight murine and primate species (mean maximum of 0.0246; Komologorov-Smirnov, K = 0.18, P = 5.26 *×* 10^-22^) (**Figure S3B**). Thus, experimentally-validated regions of the 3D genome conserved between diverse murine and primate species are also minimally 3D divergent among bonobos and chimpanzees as expected, validating our approach.

While most pairs revealed similar genome folding, many thousands had high 3D divergence (**Figure 1D**). For context, we compared the distribution of divergence scores to those generated genome-wide from pairs of 130 modern humans (Gilbertson et al., in prep). *Pan* 3D divergence is significantly higher (mean = 0.008) than the modern human distribution (mean = 0.003; Komolgorov-Smirnov, K = 0.329, P = 2.23 *×* 10^-308^) (**Figure S4**). This likely reflects the older divergence between bonobos and chimpanzees, *∼* 1.9 Ma (de Manuel et al., 2016), compared to extant human populations: *∼* 150 to 350 ka (Fan et al., 2023; Schlebusch et al., 2017). Further, greater divergence could also be explained by greater overall genetic diversity observed in *Pan* compared to modern humans, particularly among central and eastern chimpanzees (Prado-Martinez et al., 2013).

### 2.3 Genome-wide 3D divergence recapitulates *Pan* phylogeny

The distinct demographic histories among the five extant *Pan* lineages have resulted in variable genetic diversity, particularly among the four chimpanzee subspecies (Prado-Martinez et al., 2013). However, it is not known if 3D genome variation follows similar lineage-specific patterns. To investigate this, we analyzed inter-individual differences in mean 3D divergence within and among different *Pan* lineages. We first quantified this variation among all 56 individuals genome-wide by calculating the mean 3D divergence per pair.

Hierarchical clustering of mean 3D divergence per pair confirmed that 3D divergence recapitulates *Pan* phylogeny based on sequence similarity (**Figure 2A**). This clustering also emphasizes 3D divergence among individuals of different lineages. On average, interspecific pairs were the most 3D divergent, pairs comprising individuals from different chimpanzee subspecies were moderately 3D divergent, and pairs of individuals within the same lineage were the least 3D divergent (**Figure 2A**).

**Figure 2:**
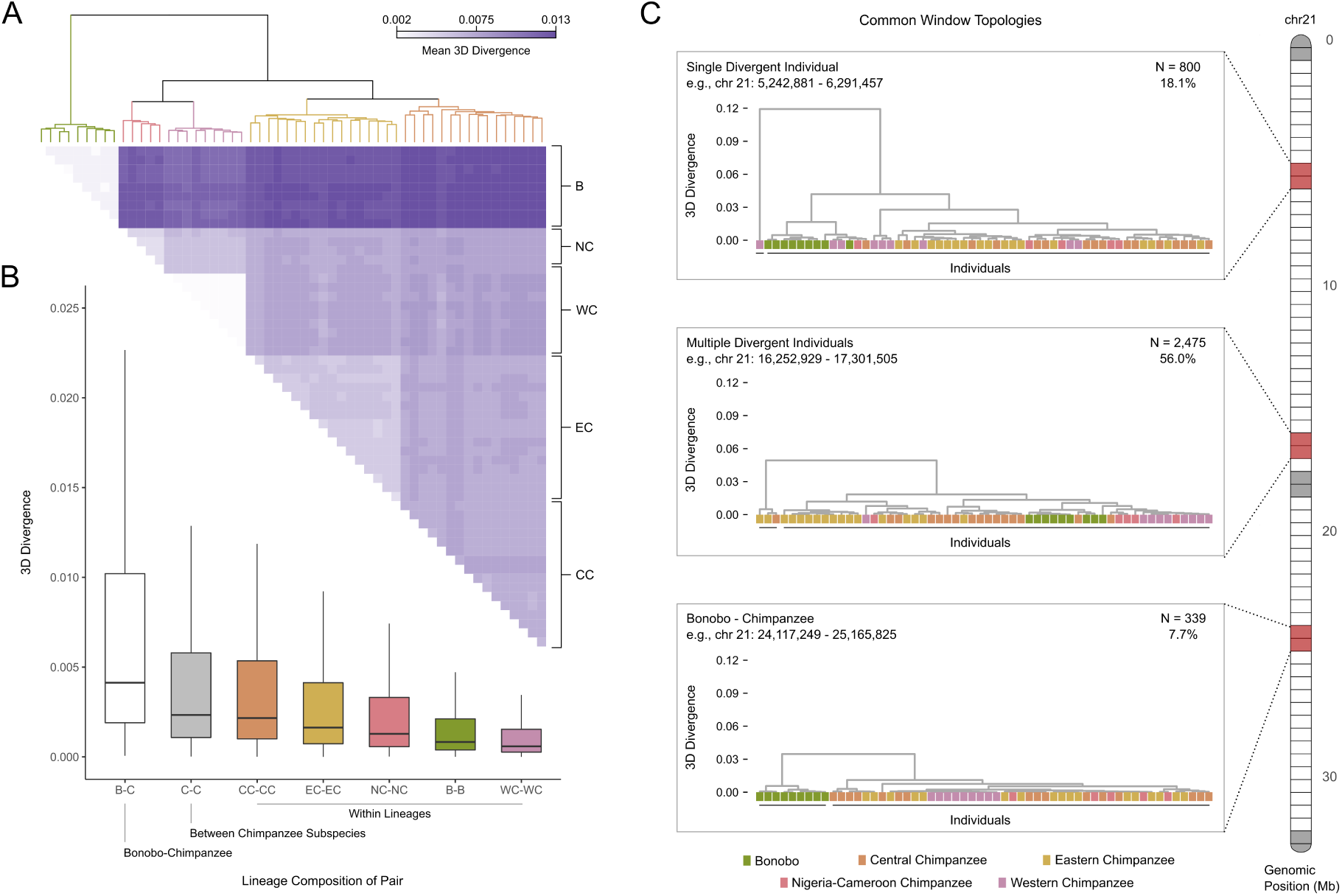
Genome-wide 3D divergence patterns recapitulate *Pan* phylogeny but are highly variable across the genome. **(A)** Mean genome-wide 3D divergence among all individuals. Rows/columns are ordered based on hierarchical clustering of 3D divergence. Lineage clusters are colored in the dendrogram and annotated on the right side of the matrix. B = bonobos, CC = central chimpanzees, EC = eastern chimpanzees, NC = Nigeria-Cameroon chimpanzees, W = western chimpanzees. **(B)** Pairwise mean genome-wide 3D divergence distributions stratified by the lineages of the individuals in each comparison. Bonobo-chimpanzee pairs and pairs of chimpanzees from different subspecies have higher 3D divergence than within lineage pairs. **(C)** Representative examples from chromosome 21 of the most common 3D divergence patterns across the genome. We hierarchically clustered all individuals based on their pairwise divergence patterns for each genomic window and found substantial variation. Here, we highlight the three most common topologies using example windows from chromosome 21: 1) a highly divergent individual, 2) multiple divergent individuals, and 3) bonobo-chimpanzee clustering. 800 (18.1%) windows are characterized by a single individual whose 3D contact pattern was an outlier compared to all others. The most common pattern (N = 2,475) consisted of multiple divergent individuals representing a subset from a single or multiple lineages. We also identified 339 windows where bonobos and chimpanzees formed separate clusters. Clusters are indicated by a black line under the individuals. Each example’s genomic position is indicated by the red shaded windows on the chromosome. Light grey shaded cells are windows that were not analyzed in this study due to missing reference sequence. In addition to the patterns illustrated here, there are eight two-cluster windows where western chimpanzees formed a lineage-specific cluster and 798 windows with *≥* 3 clusters.

The 3D divergence observed between central chimpanzees pairs (median = 0.00216) nearly encompasses the variation observed in pairs of chimpanzees from different subspecies (median = 0.00233) (**Figure 2B**). This likely reflects the high sequence diversity in central chimpanzees, which is greater than any other *Pan* lineage (Prado-Martinez et al., 2013). We also observed that median 3D divergence for within lineage pairs was positively associated with effective population size (**Table S1**).

### 2.4 3D divergence varies across the genome

While genome-wide patterns characterize variation in genome folding among individuals and lineages overall, levels of 3D divergence likely vary across the *Pan* genome. To explore this, we clustered all individuals based on 3D divergence in each of the 4,420 genomic windows separately. This approach yielded between two and five clusters per window (**Table S2**), of which two cluster windows were by far the most common (81.9%). Next, we distinguished the topologies of two cluster windows based on two characteristics: 1) the number of individuals per cluster and 2) the lineages present in each cluster (**File S2**). We identified three common topologies among the two cluster windows (**Figure 2C**).

The most common were 2,475 or 56% of windows with two clusters both comprised of multiple individuals, where the smaller cluster contained a subset of, but not all, individuals from one or more lineages. We refer to this topology as “multiple divergent individuals” clustering. To better understand these windows, we quantified the size of the smaller cluster. These clusters ranged in size from two to 28 individuals and had a median size of seven (**Figure S5A**). However, many clusters containing the divergent individuals were small; 28.3% of windows had a cluster size of 2 or 3 individuals. We also examined the lineage composition of these clusters and predicted that many would include a subset of 1) central chimpanzees, 2) eastern chimpanzees, or 3) both due to the high genetic diversity and larger sample sizes from those lineages. Eastern and central chimpanzees are the most recently diverged among *Pan* lineages and share many polymorphisms. Indeed, the most frequent lineage composition of these clusters were both eastern and central chimpanzees (N = 382), followed by central chimpanzees (N = 252), and bonobos (N = 240) (**Figure S5B**). These observations implicate the occurrence of variants present in more than one individual that result in non-lineage-specific patterns of 3D divergence in these windows.

The second most prevalent were 800 or 18.1% of windows, characterized by a single divergent individual that was assigned to its own cluster and all others to a second— i.e., “single divergent individual” clustering. We first evaluated whether these windows were the result of one or a few individuals that were frequently divergent to all others. We quantified the number of windows in which each individual was the divergent individual. All 56 individuals were the divergent individual at least once, and the frequency ranged from 1 to 34 (**Figure S6A**). Thus, these patterns are not restricted to specific individuals and are, in fact, common. Beyond frequency, the degree of 3D divergence between a divergent individual and the others varied. We retrieved the maximum 3D divergence for all windows with this topology (N = 800) and calculated the minimum, mean, and maximum 3D divergence for each individual’s set of windows (**Figure S6B**). The minima of these distributions was consistently low, suggesting that some windows may not yield consequential differences from genome folding. Distribution means were also largely similar, except for one western chimpanzee whose mean 3D divergence maximum was 0.32. As expected, distribution maxima were the most variable. 50% of individuals had a maximum 3D divergence > 0.25 (**Figure S6B**), suggesting that many of these rare divergent 3D contact patterns could have functional effects. There was no discernible pattern in frequency or distribution maxima when stratifying by lineage.

Third most common, we identified 339 or 7.7% of windows where all bonobos and chimpanzees clustered separately, i.e., “bonobo-chimpanzee” clustering. Windows in these three common topologies were significantly different in their distributions of maximum 3D divergence. Single divergent individual clustering windows had the highest mean divergence (0.067), followed by multiple divergent individuals (0.053), and bonobo-chimpanzee (0.049) (Kruskal-Wallis, H = 31.1, P = 1.77 *×* 10^-7^) (**Figure S7**). In addition to these common topologies, we also searched for other windows exhibiting lineage-specific patterns among chimpanzee subspecies. We found eight where western chimpanzees clustered separately from all other individuals (**Figure S8**, **Table S3**). Yet, we found no such windows for central, eastern, or Nigeria-Cameroon chimpanzees.

### 2.5 Interspecific 3D genome folding highlights candidates for species-specific phenotypes

The 339 genomic windows where bonobos and chimpanzees cluster separately based on 3D divergence may be evolutionarily relevant and underlie phenotypic divergence between these species. These windows spanned all analyzed chromosomes and composed 252 distinct loci after merging overlapping divergent windows. We observed striking differences when comparing bonobo and chimpanzee contact maps among many of these windows, including contact differences at binding sites for CTCF—a transcription factor and critical determinant of 3D genome structure. We identified CTCF peaks using data generated from chimpanzee LCLs (Schwalie et al., 2013). For example, interspecific 3D divergence at chr5: 16,252,929–17,301,505 ranged from 0.0249 to 0.0367 and is driven by the presence of a chimpanzee-specific “architectural stripe” that is absent in bonobos (**Figure 3A**). While both species share contact among many loci between between 16.85 and 17.1 Mb, including a CTCF peak and the *MYO10* promoter, the chimpanzee-specific stripe connects additional loci, including an upstream CTCF peak and the *MYO10* promoter. *MYO10* is a member of the myosin gene superfamily, which encode actin-based motor proteins (Berg et al., 2000). This gene is broadly expressed and knockout experiments highlight its role in many aspects of mammalian development, including the neural tube (Heimsath et al., 2017). Among adult bonobos and chimpanzees, chimpanzees exhibit higher kidney *MYO10* mRNA expression than bonobos (**Figure 3A**) (Brawand et al., 2011); however, levels are similar between species in cerebellum, heart, and liver tissue (**Figure S9**). The three other genes in this window also exhibit species differences in expression for at least one tissue (**Figure S9**). Both *ZNF622* and *RETREG1*, which are on the same strand as *MYO10* and appear to be affected by the same bonobo architectural stripe (**Figure 3A**), also have greater kidney expression in chimpanzees than bonobos.

**Figure 3:**
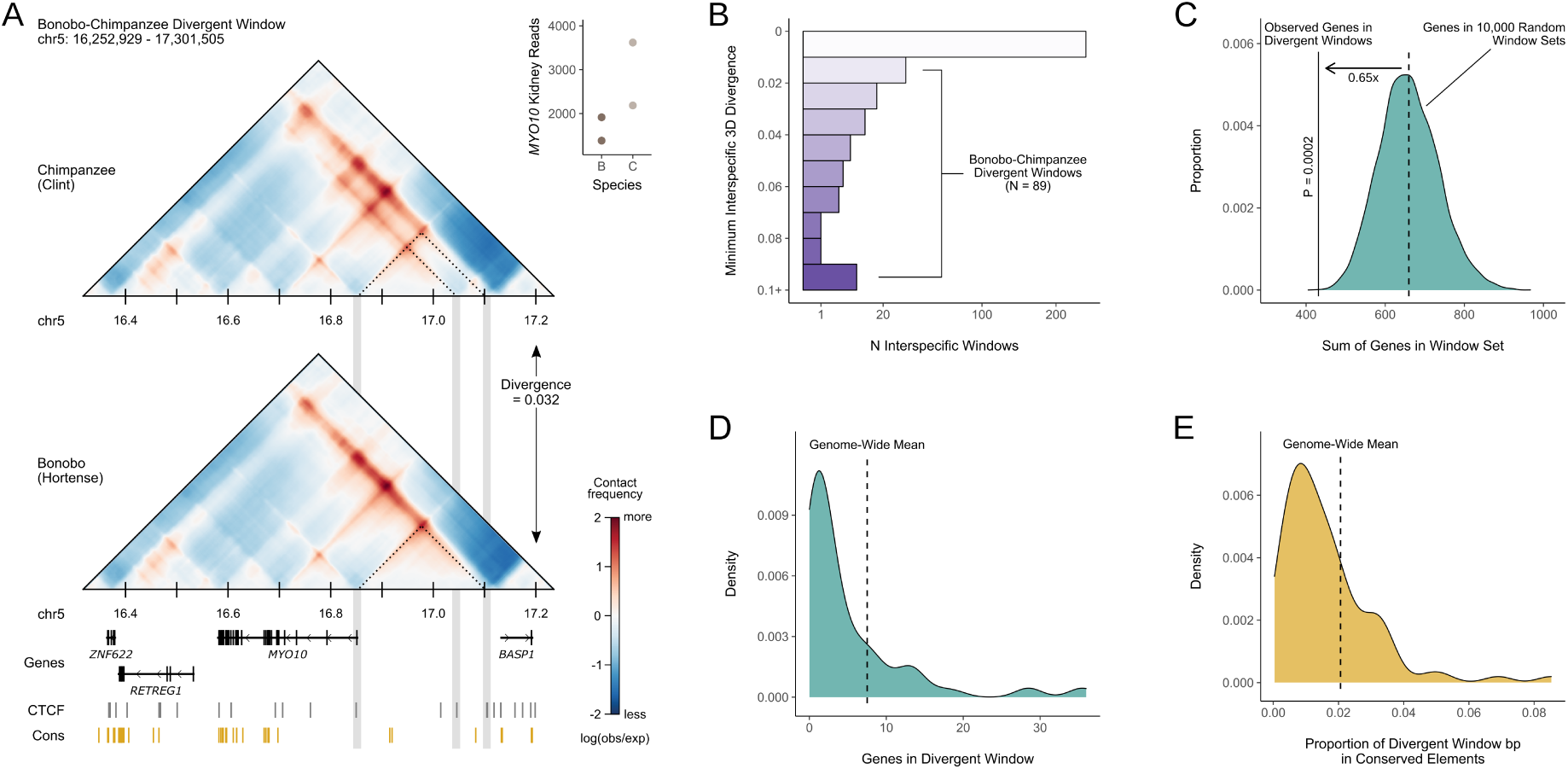
89 genomic windows have high 3D divergence between bonobos and chimpanzees. **(A)** 3D contact maps for a chimpanzee (Clint) and a bonobo (Hortense) at a representative bonobo-chimpanzee divergent window. A chimpanzee-specific “architectural stripe” indicates increased contact between many loci, including a CTCF site with the *MYO10* promoter. Dotted lines and grey boxes highlight this contact, as well as a contact between the *MYO10* promoter and another downstream CTCF site present in both species. CTCF peaks are from chimpanzee LCLs (Schwalie et al., 2013), and conserved elements (LOD > 500) are the vertebrate 30-way phastCons elements from the UCSC Genome Browser. *MYO10* read counts in kidney tissue are also shown for two bonobo (B) and two chimpanzee (C) samples from Brawand et al., 2011. **(B)** Minimum interspecific 3D divergence among 339 windows for which bonobos and chimpanzees cluster. We defined bonobo-chimpanzee divergent windows as those with a two cluster topology that completely partitions the species and a minimum interspecific divergence *≥* 0.01 (**Figure S10**). Note the x-axis is square root transformed. **(C)** Comparison of the observed vs. expected number of genes among all bonobo-chimpanzee divergent windows. We summed all genes present in the 89 windows, removing duplicates. We generated a null distribution by permuting 89 windows among the 4,420 analyzed windows 10,000 times and counting genes as for the observed set. The observed bonobo-chimpanzee divergent windows are significantly depleted of genes (0.65x expected, P = 0.002, one-tailed permutation test). The null distribution ranged from 418 to 967 genes, with a mean of 659.68. **(D)** The distribution of observed gene counts per window among bonobo-chimpanzee divergent windows. The dashed line indicates the genome-wide mean: 7.51. **(E)** The distribution of observed conserved element proportions per window among bonobo-chimpanzee divergent windows. Proportions were calculated as the sum of bp in a given window overlapping primate phastCons elements divided by the window length: 1,048,576 bp. The dashed line indicates the genome-wide mean: 0.021.

We focused on 89 “bonobo-chimpanzee divergent” windows with large and consistent interspecific 3D genome divergence (minimum 3D divergence *≥* 0.01; **Figures 3B**, **S10**). Bonobo-chimpanzee divergent windows exhibited multiple striking characteristics. First, they are significantly depleted of genes (**Figure 3C**; P = 0.002, one-tailed permutation test), and they include 17 windows with zero genes (**Figure 3D**). 431 genes unique genes are found in these windows; this is 0.65x the expected gene overlap if divergent windows were randomly distributed across the analyzable genome. Bonobo-chimpanzee divergent windows also exhibit less sequencelevel constraint between species than expected from the genome-wide distribution (**Figure 3E**). We quantified constraint as the fraction of bp in each window found in phastCons conserved elements called on a 30-way alignment of vertebrates, and 74% of windows (N = 66) were below the genome-wide conserved element fraction. However, for both gene density and conserved elements, we also observed a second set of divergent windows with higher density than expected from the genome-wide distribution (**Figures 3D**, **3E**). For example, 20 windows had *≥* eight genes, including a high-density window overlapping 36 genes. Taken together, these characteristics indicate that most species-specific genome folding occurs in genomic regions with weak evolutionary constraint and few functional elements. However, species-specific patterns also occur in a smaller number of regions that have more constraint and functional elements, and thus are more likely to contribute to changes between species.

We also explored whether genes in bonobo-chimpanzee divergent windows were enriched for genes associated with annotated phenotypes, particularly those known to differ between species. We considered annotations from the 2021 Biological Process Gene Ontology (Ashburner et al., 2000; The Gene Ontology Consortium, 2021), the 2019 GWAS Catalog (Buniello et al., 2019), the Human Phenotype Ontology (HPO; Köhler et al., 2021), and the Level 4 2021 MGI Mammalian Phenotype Ontology (MP; Eppig et al., 2015; Smith and Eppig, 2009). We did not identify any enriched traits at FDR-adjusted significance-levels (**Figure S11**, **File S3**). However, we noted a handful of phenotypes with modest enrichment related to traits that differentiate bonobos and chimpanzees including abnormality of the labia major (HPO, enrichment = 3.85, P = 0.09) and decreased body mass index (MP, enrichment = 4.32, P = 0.03) (**File S3**).

### 2.6 Individual variants drive species-specific genome folding

Next, to better understand the determinants of species-specific genome folding, we investigated sequence differences among the bonobo-chimpanzee 3D divergent loci. We quantified the contribution of different alleles to predicted 3D genome divergence using *in silico* mutagenesis (Gunsalus et al., 2023a; McArthur et al., 2022). First, we identified all bonobo-specific variants among bonobo-chimpanzee divergent windows, i.e., sites where all nine bonobos analyzed were heterozygous or homozygous for the non-reference allele and all chimpanzees were fixed for the reference allele (**Figure 4A**). We identified 115,191 total variants and 127,075 variant-window pairs, as some variants are present in overlapping divergent windows. Next, we inserted each bonobo-specific variant into the chimpanzee reference sequence for the window, predicted chromatin contacts using Akita, and calculated the 3D divergence between the full chimpanzee reference sequence and the reference with each variant (**Figure 4A**). Variants were defined as “3D-modifying” if the resulting 3D divergence between reference and mutated reference was *≥* the minimum 3D divergence score among bonobo-chimpanzee pairs for that window. We also applied this approach to lineage-specific variants among the four chimpanzee subspecies (**Supplementary Information**).

**Figure 4:**
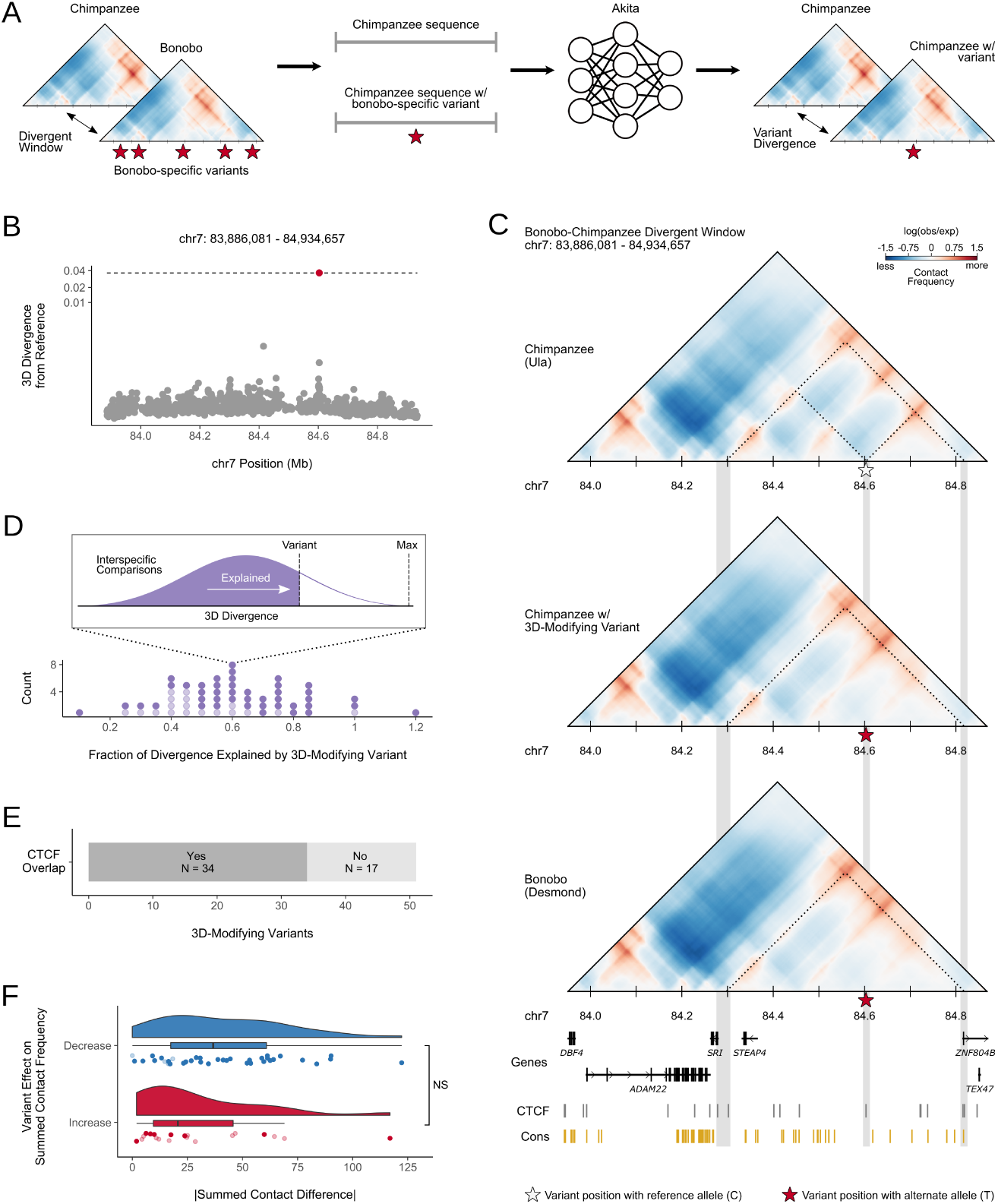
*In silico* mutagenesis reveals 3D-modifying variants that contribute to species-specific 3D genome folding patterns. **(A)** Schematic of using *in silico* mutagenesis to identify SNVs that contribute to 3D genome differences between bonobos and chimpanzees. This procedure identified 61 variant-window pairs, which consisted of 51 unique variants. **(B)** 3D divergence from reference sequence for 1,425 bonobo-specific variants within the chr7: 83,886,081–84,934,657 window. Only one variant (red dot), chr7: 84,603,122 (C > T), results in 3D divergence (0.0367) that is *≥* the observed minimum divergence among bonobo-chimpanzee pairs for this window (dashed line) (0.0366). Note that the y-axis is cube root transformed. **(C)** Contact maps for a chimpanzee (Ula), panTro6 sequence with a 3D-modifying variant, and a bonobo (Desmond) at a bonobo-chimpanzee divergent window (chr7: 83,886,081– 84,934,657). A bonobo-specific 3D-modifying variant at chr7: 84,603,122 (C > T) reduces contact between a CTCF peak and the promoters for *SRI* and *ZNF804B* compared to chimpanzees. Insertion of this variant into chimpanzee sequence recapitulates bonobo genome folding at this window. The position of this variant is indicated by a star and colored based on the input allele for the contact map (C = grey, T = red). Dotted lines and grey bars highlight relevant contacts and annotations associated with the 3D-modifying variant. **(D)** The distribution of divergence explained by the 61 3D-modifying variant-window pairs. We calculated explained divergence by dividing the variant divergence score by the maximum interspecific divergence observed for a given window. Explained divergence counts are displayed in 0.05 bins. CTCF overlap is indicated by shading (light = no overlap, dark = overlap). **(E)** The number of 3D-modifying variants that fall within and outside CTCF peaks identified using chimpanzee LCLs (Schwalie et al., 2013). **(F)** Summed contact differences induced by 3D-modifying variants stratified by net effect. We summed all contact frequencies with the 2,048 bp bin containing the 3D modifying variant for the reference map and reference with 3D-modifying variant map (**Figure S14**). The contact difference was calculated by subtracting the reference contact sum from the reference with variant contact sum. Thus, positive values indicate increased contact overall due to the 3D-modifying variant, while negative values indicate decreased contact overall. We used the absolute values of summed contact differences to compare 3D-modifying variants that increase and decrease contact overall. These distributions were not significantly different (Mann Whitney U, U = 483, P = 0.09). Individual variant effects are indicated by points and distributions are illustrated with box and violin plots. Color indicates overall effect. CTCF overlap is indicated by shading (light = no overlap, dark = overlap).

The interspecific 3D divergence among 59 (66.3%) of the bonobo-chimpanzee divergent windows could largely be recapitulated by inserting a single variant. For example, among the 1,425 variants intersecting the genomic window at chr7: 83,886,081–84,934,657, only one variant resulted in substantial 3D divergence (**Figure 4B**). Chimpanzees are fixed for the C allele at chr7: 84,603,122, while all bonobos have at least one T allele. The bonobo allele appears to result in decreased contact with promoters for *SRI* and *ZNF804B* and increased contact among loci adjacent to the variant (**Figure 4C**). *SRI* is a penta-EF hand calcium binding protein, regulating intracellular calcium and mediating excitation-contraction coupling in heart and skeletal muscle (Meyers et al., 1998), and has been implicated in neurodegenerative disease (Mattson et al., 2000). The function of *ZNF804B* is largely unknown; however, this gene is largely expressed in thyroid tissue among human adults (https://www.proteinatlas.org/ENSG00000182348-ZNF804B/tissue#rna_expression). This observation is intriguing because bonobos and chimpanzees developmentally differ in thyroid levels (Behringer et al., 2014). *ZNF804* also exhibits a species difference in cerebellum expression, whereas *SRI* and two other nearby genes (*STEAP4*, *TEX47*) do not (**Figure S12**).

Interspecific 3D divergence at this window ranges from 0.0366 to 0.07. When inserted into the chimpanzee reference sequence, the T allele resulted in 3D divergence of 0.0367 from the reference sequence (**Figure 4C**). Therefore, this variant appears to drive most of species difference observed at this locus. However, other bonobo-specific variants likely explain additional divergence in the interspecific comparison distribution.

Overall, 51 bonobo-specific variants were 3D-modifying, of which ten occurred among overlapping divergent windows, resulting in 61 3D-modifying variant-window pairs (**File S4**). Two windows also contained two separate 3D-modifying variants—chr4: 113,770,497–114,819,073 and chr10: 87,556,097–88,604,673. These variants were four and two nucleotides apart, respectively, suggesting the perturbation of the same genomic element. We predicted that most 3D-modifying variants are derived alleles. We tested this hypothesis by comparing the 3D-modifying allele and the inferred ancestral allele to quantify the proportion of ancestral and derived variants. Ancestral allele calls were determined using a probabilistic method to infer ancestral sequence from multiple primate sequences (Martin et al., 2023). Ten 3D-modifying variants were ancestral, whereas 41 were derived.

We quantified the fraction of the observed bonobo-chimpanzee divergence a given variant “explained” by dividing the 3D divergence from *in silico* mutagenesis by the observed interspecific maximum (**Figure 4D**). For example, the aforementioned variant at chr7: 84,603,122 explained 52% of the maximum interspecific divergence observed for its window (chr7: 83,886,081– 84,934,657). Surprisingly, the 3D-modifying variants often explained a considerable fraction of 3D divergence (mean = 0.57). Three variants explained approximately 100% of the divergence observed in their windows and one explained 117% of the observed divergence suggesting that other variants in the window likely buffer against the variant’s 3D-modifying effect. Conversely, the presence of multiple variants with small to modest effects may also result in species-specific genome folding patterns. This hypothesis may explain windows where no 3D-modifying variants were identified or those windows with variants that minimally explained divergence. Thus, differences in genome folding between bonobos and chimpanzees are largely driven by individual variants with large effects, yet other differences may occur due to multiple variants with small effects.

### 2.7 CTCF binding motif disruption explains many, but not all of the bonobo-chimpanzee 3D divergent windows

We anticipated that many 3D-modifying variants would fall within the binding domains of CTCF, as in the example window (**Figure 4C**). Indeed, 34 (66.67%) of 3D-modifying variants intersected CTCF peaks (**Figure 4E**). Two additional 3D-modifying variants fell within 10 kb of a CTCF peak. 3D-modifying variants overlapping CTCF peaks explained significantly more 3D divergence (mean = 0.61) than those that did not (mean = 0.47) (Mann-Whitney, U = 207, P = 0.005). Next, we quantified the mutation spectrum of the 3D-modifying variants; we were particularly interested to see if C > T mutations promoted by GC-biased gene conversion at CpG sites were common. 14 (27.5%) of the variants were C > T mutations; however, these were not enriched for CpGs (**Figure S13**). This suggests that 3D-modifying variants that contribute to species differences in genome folding are unlikely to be the result of GC-biased gene conversion and are largely, but not entirely, driven by mutations that modify CTCF binding motifs. We also quantified the effects of 3D-modifying variants on contact frequency. For example, the chr7: 84,603,122 variant results in decreased contact between that locus and other loci in the window. We predicted that most 3D-modifying variants would similarly decrease contacts, because we anticipated that derived variants are more likely to disrupt functional motifs, e.g., for CTCF or other transcription factors, than to create a new functional element. We classified each variant-window pair as resulting in decreased or increased overall contact for the variant locus by subtracting the summed values from all cells in the contact map representing contacts with the variant locus (N = 448) between the variant map and reference map (**Figure S14**). Thus, positive values indicate increased contact overall due to the 3D-modifying variant, while negative value indicate decreased contact overall. As predicted, 3D-modifying variants more frequently result in decreased (N = 38) rather than increased contact (N = 20) (**Figure 4F**).

When stratified by CTCF overlap, 8 or 40% of variants that increased chromatin contact overall fell within a CTCF peak, while 34 or 89.5% of variants resulting in decreased chromatin contact overlapped a CTCF peak. We also stratified chromatin contact effects by allele age and found that ancestral and derived variants occurred in similar proportions among variants that increased contact, 44% and 56%, respectively. However, derived variants comprised the majority (90%) of variants resulting in overall decreased contact. These patterns broadly support the hypothesis that 3D-modifying variants are more likely to disrupt CTCF binding sites, resulting in decreased contact. Conversely, it also appears that variants outside of CTCF sites, perhaps overlapping other transcription factors, often yield increased chromatin contact. We used the absolute values of summed contact differences to compare variants that increased vs decreased contact and did not find a significant difference between these distributions (Mann-Whitney, U = 483, P = 0.09) (**Figure 4F**). Therefore, 3D-modifying variants in *Pan* are more likely to result in decreased chromatin contact via CTCF disruption but the measurable effect is comparable between variants that overall decrease or increase contact.

## 3 Discussion

The complex 3D organization of the nuclear genome plays an important role in cell biology, particularly gene regulation, and disruption of genome folding is associated with phenotypic variation and disease in humans and other species (Lupiáñez et al., 2015; Norton and Phillips-Cremins, 2017). These observations have prompted close examination of 3D genome variation both within and among diverse species using experimental data (Dixon et al., 2012; Eres et al., 2019; Li et al., 2022; Li et al., 2023; Lukyanchikova et al., 2022; Torosin et al., 2022; Yang et al., 2019). While 3D genome data are available for humans and other model organisms, data remain scarce for other species. Further, generation of genome folding data at remains challenging to accomplish at population-scale. The development of machine learning algorithms (Fudenberg et al., 2020; Schwessinger et al., 2020; Zhou, 2022) that learn from existing data to predict 3D genome folding from sequence alone offer an opportunity to close this knowledge gap. Here, we apply machine learning methods to rapidly assay variation in genome folding in humans’ closest living relatives.

Much of the inferred *Pan* 3D genome is similar among all five extant lineages, including conserved TAD boundaries identified from experimental data. However, a small fraction of the genome displays substantial variation in chromatin contact. Genome-wide patterns of 3D divergence recapitulate the *Pan* phylogeny; yet, individual genomic windows harbored more complex patterns, including many windows characterized by a single or several individuals with divergent chromatin contact patterns. We identify loci characterized by species-specific genome folding that contain different contact patterns that co-localize with gene expression differences between species.

Our computational approach enables the rapid prediction of genome folding from DNA sequence alone. The ability to rapidly scan the effects of candidate variants enables prioritization of variants and loci for experimental validation studies. Applying this *in silico* mutagenesis approach to *Pan*, we identify variants that likely contribute most to species differences in genome folding. We find that the patterns at many divergent windows are driven by a single SNV that disrupts CTCF binding. These findings reveal the potential of genome folding at specific loci to contribute to phenotypic divergence in humans’ closest living relatives.

Non-coding variation comprises the majority of genetic variation in *Pan*; however, the consequences and the specific mechanisms through which non-coding variants regulate gene expression remain largely unknown in these taxa. We illuminate one of these mechanisms here and propose that some gene expression differences are associated with 3D genome variation between bonobos and chimpanzees. Our work also contributes to a broader context to comparisons of chromatin contact at population-scale in recent primate evolution. For example, we observed considerably higher 3D divergence in *Pan* than between archaic hominins and modern humans as well as within modern humans (Gilbertson et al., in prep; McArthur et al., 2022).

This research represents an important first step in understanding *Pan* 3D genome variation; however, we recognize the limitations of the present study and the promise of future research. First, the expansion of available data and development of new algorithms may yield more accurate models for predicting the 3D genome from sequence. Such advances may enable predictions at higher resolution, incorporation of other variant types (e.g., structural variants), and for specific tissue and cellular contexts (Tan et al., 2023; Zhou, 2022). Second, our ability to fully understand the functional consequences of differences in chromatin contact is limited by the currently available functional annotations. Additional data on transcription factor binding and RNA across tissues and cells in these species will help fully realize species differences in genome folding and benefit the study of other regulatory mechanisms.

In conclusion, we demonstrate utility of applying DNA sequence-based machine learning to the genomes of non-model systems that lack the rich functional and experimental data available for humans. Our findings shed light on an important gene regulatory mechanism in humans’ closest living relatives and identify loci that may contribute to phenotypic divergence in *Pan*.

## 4 Methods

### 4.1 *Pan* Genomic Data

We retrieved raw short read data from the Great Ape Genome Project (de Manuel et al., 2016; Prado-Martinez et al., 2013), representing high-coverage genomes from 13 bonobos (*Pan paniscus*), 18 central chimpanzees (*P. troglodytes troglodytes*), 19 eastern chimpanzees (*P. t. schweinfurthii*), 10 Nigeria–Cameroon chimpanzees (*P. t. ellioti*), and 11 western chimpanzees (*P. t. verus*).

We used genotypes generated in Brand et al., 2021. Briefly, we mapped short reads to a current high-quality chimpanzee reference genome, panTro6 (Kronenberg et al., 2018), using sex-specific versions of the reference generated from XYAlign (Webster et al., 2019). We used bcftools, version 1.18 (Li, 2011) to filter genotypes. We included high-quality sites with biallelic SNVs where all 71 genotypes were called. We chose to exclude structural variants due to the difficulty of classifying chromtain contact among sequences of different lengths. Next, we set low-quality genotypes to the reference allele and excluded sites that were fixed for the reference allele.

The number of variants among individuals from each *Pan* lineage (**File S1**, **Figure S2**) were consistent with phylogenetic predictions as the reference sequence is a western chimpanzee (bonobos > eastern/central chimpanzees > Nigeria-Cameroon/western chimpanzees). We observed a handful of individuals per lineage with substantially fewer SNVs compared to others from the same lineage. Most of this variation appears to be driven by low quality genotypes that did not pass filtering. We excluded these individuals (N = 15) from downstream analyses (**File S1**, **Figure S2**).

We generated pseudo-haploid sequences for each individual using GATK’s FastaAlternateReferenceMaker (Poplin et al., 2018) to add the quality-filtered SNVs to the reference sequence. This approach considers heterozygotes and homozygotes for the non-reference allele to be equivalent. We excluded unlocalized scaffolds (N = 4), unplaced contigs (N = 4,316), the Y chromosome, and mitochondrial genome from these sequences.

### 4.2 3D Genome Predictions with Akita and Model Performance on Chimpanzee Sequence

We used a convolutional neural network, Akita, to predict 3D genome organization from the pseudo-haploid sequences (Fudenberg et al., 2020). A detailed description of the CNN can be found in Fudenberg et al., 2020. Briefly, Akita uses an input sequence of length 1,048,576 bp to output predicted chromatin contact for the central 917,504 bp of the input sequence at 2,048 bp resolution. Each cell value is log2(obs/exp)-scaled because chromatin contact is distance dependent. The Hi-C maps used to train Akita were clipped to contact frequencies between -2 and 2 (Fudenberg et al., 2020). Thus, as expected, most predicted values range from -2 to 2 (**Figure S15**). Akita was simultaneously trained on five cell types from Hi-C and Micro-C datasets: GM12878, H1ESC, HCT116, HFF, and IMR90 (Fudenberg et al., 2020).

Before we applied Akita to DNA sequences of different *Pan* lineages, we evaluated the accuracy of Akita on chimpanzee sequences by comparing the predictions with the experimental Hi-C data. Briefly, we lifted over the regions in the human test set from hg38 to panTro6 using liftOver (Hinrichs et al., 2006), retaining regions of window size within +/- 10% of 1,048,576 bp and with less than 1% of missingness, and extracted their DNA sequences as input to Akita. Of the outputs in five different cell types, we focused on the predictions for human foreskin fibroblast (HFF) following McArthur et al., 2022. The Hi-C data were obtained from chimpanzee neural progenitor cells (NPC) (Keough et al., 2022), rebinned into 2,048-bp bins using cooler (Abdennur and Mirny, 2020) and then processed as previously described for human datasets in Fudenberg et al., 2020.

### 4.3 Chromatin Contact Map Generation and Comparison

We segmented the panTro6 reference sequence by creating a sliding 1,048,576 bp window per chromosome that overlapped by half, resulting in 5,317 total windows. We discarded windows without complete sequence coverage (i.e., *≥* 1 “N”s), including any incomplete windows at the end of each chromosome, retaining 4,420 windows.

We used Akita to create 3D genome predictions from the pseudo-haploid sequences per window per individual. We output predictions for both HFF and GM12878 and compared all autosomal windows between all pairs of individuals (N = 6,541,920) as well as X chromosome windows between all pairs of females (N = 95,130) because that chromosome is hemizygous in males. Comparisons were made by calculating the mean squared error and Spearman’s *ρ* between a pair of contact maps. Next, we calculated a third metric from the latter, “3D divergence” (1 - *ρ*). Lower 3D divergence reflects similarity between a pair of contact maps whereas higher 3D divergence indicates differences between a pair of maps.

We contrasted the resulting distribution of *Pan* 3D divergence to a distribution generated from 130 modern humans (Gilbertson et al., in prep). Five individuals were sampled from each of the 26 subpopulations from the Thousand Genomes Project (Auton et al., 2015). Contact maps and pairwise 3D divergence were generated as above for all autosomal windows without missing coverage in the hg38 reference assembly, resulting in 40,860,105 total comparisons. We compared the *Pan* and modern human distributions using a Komologorov-Smirnov test.

### 4.4 3D Divergence at Primate-conserved and Ultraconserved TAD Boundaries

We compared the distribution of *Pan* 3D divergence overlapping experimentally validated conserved TAD boundaries to the genome-wide distribution. We used two sets of 10 kb conserved boundaries among autosomes and the X chromosome from Okhovat et al., 2023: 1) “primate-conserved” boundaries (N = 491), defined as conserved among *Homo sapiens*, *Hylobates moloch*, *Nomascus leucogenys*, and *Macaca mulatta*, and 2) “ultraconserved” boundaries (N = 1,023), defined as conserved among all four primate species as well as four murines—*Mus caroli*, *M. musculus*, *M. pahari*, and *Rattus norvegicus*. We used liftOver (Hinrichs et al., 2006) with all default settings to convert boundaries from hg38 to panTro6 coordinates, resulting in 415 primate-conserved and 915 ultraconserved boundaries. Next, we retrieved the maximum 3D divergence for windows overlapping the primate-conserved and ultraconserved boundaries as well as the maxima for all 4,420 windows as the genome-wide set using Pybedtools intersect, version 0.9.0 (Dale et al., 2011). However, we anticipated that some TAD boundaries would occur in overlapping windows yielding two maxima per boundary. Therefore, we decided to identify the window in which the TAD boundary was most central by calculating a centrality score for each TAD boundary/window pair:

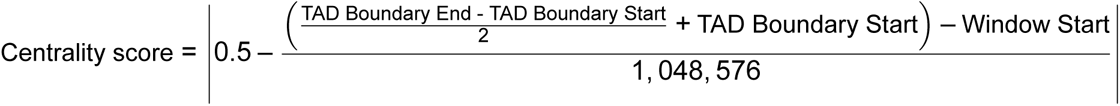

Scores at or near 0 indicate the TAD boundary is more central to a given window, whereas values closer to 0.5 indicate the TAD boundary is near the edge of a given window. We compared the distribution of 3D maxima in both these sets to the genome-wide distribution using a Komologorov-Smirnov test.

### 4.5 Hierarchical Clustering and Window Topology Analysis

We performed hierarchical clustering on the pairwise 3D divergence scores for all individuals per genomic window. Hierarchical clustering was implemented using SciPy, version 1.9.1 (Virtanen et al., 2020). We used complete linkage, which is robust to outliers and generates separate, spherical clusters. We first identified the number of clusters per window. We further considered the size and lineage composition of each cluster among the two cluster windows. Using these characteristics, we designated three topologies for two cluster windows: 1) windows characterized by a single divergent individual that was assigned to it’s own cluster and all others to another, i.e., single divergent individual, 2) windows with clusters comprised of multiple individuals, where neither cluster was lineage-specific, i.e., multiple divergent individuals, and 3) windows with a lineage-specific cluster and another containing all other individuals. Among these lineage-specific clusters, 339 were bonobo-specific and eight were western chimpanzee-specific. We did not further characterize topologies for windows featuring three, four, or five clusters.

### 4.6 Phenotype Enrichment

We used our previous approach applying a permutation-based empirical null distribution to quantify gene enrichment in different phenotypes from a set of genomic features (McArthur et al., 2022; Brand et al., 2023). Annotations were retrieved from Enrichr (Chen et al., 2013; Kuleshov et al., 2016; Xie et al., 2021) for four ontologies: 1) 2021 Gene Ontology Biological Process, 2) 2019 GWAS Catalog, 3) Human Phenotype Ontology, and 4) 2021 MGI Mammalian Phenotype Ontology Level 4.

The Biological Process Gene Ontology (BP) domain considers annotations for processes accomplished by multiple molecular activities and the 2021 catalog includes 6,036 terms and 14,937 genes (Ashburner et al., 2000; The Gene Ontology Consortium, 2021). The 2019 GWAS Catalog (GWAS) largely considers common disease annotations and has 1,737 terms with 19,378 genes (Buniello et al., 2019). The Human Phenotype Ontology (HPO) considers rare disease annotations and has 1,779 terms with 3,096 genes (Köhler et al., 2021). The MGI Mammalian Phenotype Ontology (MP) was developed for mouse phenotypes and the 2021 Level 4 catalog includes 4,601 terms and 9,767 genes (Eppig et al., 2015; Smith and Eppig, 2009).

We identified the number of genes represented per term among the 431 genes in bonobo-chimpanzee divergent windows for each ontology, excluding terms with no representation. This resulted in the consideration of 2,135 terms from BP, 552 terms from GWAS, 621 terms from HPO, and 1,740 terms from MP.

Next, we shuffled the 89 windows randomly among all 4,420 genomic windows used in this analysis and summed the genes observed for each phenotype annotation. We repeated this process 1*×*10^4^ times per ontology and calculated enrichment as the number of observed genes divided by the mean empirical gene count per term. p-values were calculated as the proportion of empiric counts + 1 *≥* the observed counts + 1. We adjusted our significance level due to multiple testing by correcting for the false discovery rate (FDR). We used a subset (N = 1 *×* 10^3^) of the empirical null observations and selected the highest p-value threshold that resulted in a V/R < Q where V is the mean number of expected false discoveries and R is the observed discoveries (McArthur et al., 2022). We calculated adjusted significance levels for each set for Q at both 0.05 and 0.1. This analysis was run using a Snakemake, version 7.14.0, pipeline (Köster and Rahmann, 2012).

### 4.7 Gene Expression

We identified gene expression differences between bonobos and chimpanzees using RNAseq data from Brawand et al., 2011. These data primarily consist of 76 bp long single reads per tissue per species (N = 21). Cerebellum, heart, kidney, and liver were sampled once per female and male per species. Prefrontal cortex was sampled for the chimpanzee female and both bonobo individuals. Testis was also sampled from each male per species. We did not include 202 bp paired end reads from prefrontal cortex samples (N = 6) in this analysis.

We assessed read quality using fastqc, version 0.11.9 (Andrews, 2010) and multiqc, version 1.13a (Ewels et al., 2016) and identified a number of samples with mean Phred scores *≤* 20 at the first base and 3’ tail of the read as well as two samples with *≥* 1% of sequences with adapter content. We used trimmomatic, version 0.39 (Bolger et al., 2014) to filter out adapter sequences and remove the first base and bases after the 58th base, resulting in 58 bp reads. These trimmed sequences resulted in improved Phred scores per base and minimal sequences with adapter content. We then prepared the reference sequence for mapping and mapped reads to the panTro6 genome using star, version 2.7.10a (Dobin et al., 2013). Read counts per gene were calculated using htseq, version 2.0.2 (Anders et al., 2015). This analysis was run using a Snakemake, version 7.14.0, pipeline (Köster and Rahmann, 2012).

The small number of biological replicates reduces power to detect species differences in this dataset (Schurch et al., 2016) using genome-wide approaches such as DESeq2. Therefore, we restricted consideration to genes that fell within the 89 bonobo-chimpanzee divergent windows and tissues with two replicates in both species: cerebellum, heart, kidney, and liver. We excluded any gene-tissue pairs where any of the four samples had zero reads, resulting in 1,361 gene-tissue pairs. We then looked for gene-tissues pairs where bonobo and chimpanzee read counts were non-overlapping (N = 442) (e.g., *MYO10*, (**Figure 3A**). We quantified the gene expression difference as the number of reads between the maximum value of the species with lower expression and the minimum value of the species with higher expression.

### 4.8 *In Silico* Mutagenesis

We identified individual nucleotides contributing to 3D divergence among bonobo-chimpanzee divergent windows using an *in silico* approach (**Figure 4A**). We identified “bonobo-specific” alleles among the 89 bonobo-chimpanzee divergent windows, consisting of 115,191 unique variants and 127,075 variant-window pairs, due to the presence of some variants in overlapping divergent windows. “Bonobo-specific” alleles were defined as alleles present in heterozygous or homozygous genotypes for the non-reference (chimpanzee) allele among all nine bonobos, while all 47 chimpanzees were fixed for the reference allele. We considered both heterozygous and homozygous genotypes because we used pseudo-haploid sequences to predict genome folding. For each variant-window pair, we inserted the variant into the reference sequence for that window and calculated the MSE and 3D divergence between the reference map and the reference with variant map. “3D-modifying variants” were defined as variants the resulted in 3D divergence *≥* the minimum 3D divergence score among interspecific comparisons for that window.

We calculated the effects of 3D-modifying variants by calculating two metrics per variant-window pair. First, we calculated “explained divergence” by dividing the 3D divergence for the variant by the maximum interspecific comparison for the window. Values near zero indicate that the 3D-modifying variant explains minimal divergence among the observed comparisons, while values near one indicate the variant explains most of the divergence among observed comparisons. Values greater than one indicate that variant creates more 3D divergence than observed among any interspecific comparison, suggesting that other variants may “buffer” against the variant’s effect. Second, we calculated the “summed contact difference” (**Figure S14**). This metric captures the overall effect of a 3D-modifying variant by summing the contact frequencies for all cells that represent contact between the cell containing the variant and all others (N = 448 cells). We subtracted the summed contact difference of the map for the reference sequence from the map for the reference sequence with the 3D-modifying variant. Thus, positive summed contact difference values indicate overall increased contact from the 3D-modifying variant, whereas negative values indicate overall decreased contact. We excluded three variants from this calculation that fell outside the central 917,504 bp in a genomic window predicted by Akita.

We also considered whether 3D-modifying variants were ancestral or derived using ancestral alleles called using Ortheus from an EPO multi-species primate alignment (Martin et al., 2023). We used these designations to stratify chromatin contact effects but excluded three variants that occurred in overlapping divergent windows. Two disagreed in effect (“decrease” in one window and “increase” in another), which is expected due to the limited sequence overlap in overlapping windows (50%). The third variant occurred in the middle 917,504 bp output by Akita in one window but fell outside this region in another. Therefore, we excluded these three variants from quantifying the impact of allele state on chromatin contact effect, using the 48 other 3D-modifying variants for analysis.

We also applied our *in silico* mutagenesis approach to lineage-specific variants among the four chimpanzee subspecies. Lineage-specific variants were defined as before—all individuals in the lineage of interest had a genotype with at least one non-reference allele, whereas all others were fixed for the reference allele. We considered variants in all windows identifying 78 unique variants with 150 variant-window pairs in central chimpanzees, 337 unique variants with 610 variant-window pairs in eastern chimpanzees, 34,474 unique variants with 64,657 variant-window pairs in Nigeria-Cameroon chimpanzees, and 11,993 unique variants with 22,671 variant-window pairs in western chimpanzees.

### 4.9 Genomic Annotations

We retrieved various annotations to understand the context of 3D genome differences. We used gene annotations from NCBI and retained the longest transcript for genes with multiple transcripts. We used the chimpanzee CTCF annotations from Schwalie et al., 2013. These annotations were generated from LCLs from seven primates and both human and mouse livers. We retrieved phastCons elements called using a multiple species aligment of 30 species from the UCSC Genome Browser. Ancestral alleles were identified using Ensembl release 110 (Martin et al., 2023). Genomic coordinates for these annotations were converted to panTro6 using liftOver (Hinrichs et al., 2006) with all default settings.

### 4.10 Analysis

All data analyses were performed using Bash and Python scripts, some of which were implemented in Jupyter notebooks. All reported p-values are two-tailed, unless noted otherwise. The machine used to run analyses had a minimum value for representing floating numbers of 2.2250738585072014 *×* 10^308^. Therefore, we abbreviate values less than this as 2.23 *×* 10^308^.

### 4.11 Visualization

Results were visualized using Inkscape, version 1.1 (Inkscape Project, 2020) and ggplot, version 3.3.6 (Wickham, 2016) implemented in R, version 4.0.5 (R Core Team, 2020).

## 4.12 Data Availability

We used publicly available data for all analyses. The raw *Pan* data were retrieved from the Sequence Read Archive (accession nos. PRJNA189439 and SRP018689) and the European Nucleotide Archive (accession no. PRJEB15086) (de Manuel et al., 2016; Prado-Martinez et al., 2013). Ancestral alleles were retrieved from Ensembl(http://ftp.ensembl.org/pub/release-110/fasta/ancestral_alleles/homo_sapiens_ancestor_GRCh38.tar.gz). CTCF data were retrieved from the Functional Genomics Data Collection (https://www.ebi.ac.uk/arrayexpress/files/E-MTAB-1511/E-MTAB-1511.additional.1.zip). Gene expression data were retrieved from the SRA (GEO accession nos. GSM752664-GSM752690). phastCons elements were retrieved from the UCSC Genome Browser (https://hgdownload.soe.ucsc.edu/goldenPath/hg38/database/phastConsElements30way.txt.gz). The HFF pairwise comparisons file used in this analysis is available on Dryad (DOI:10.5061/dryad.7pvmcvf11).

## 4.13 Code Availability

All code used to conduct analyses and generate figures is publicly available on GitHub (https://github.com/brandcm/Pan_3d_Genome). Akita is available from the basenji repository on GitHub (https://github.com/calico/basenji/tree/master/manuscripts/akita). The pipeline used to generate the VCFs is also available on GitHub (https://github.com/thw17/Pan_reassembly).

## 4.14 Acknowledgements

We thank Mariam Ohkovat and Lucia Carbone for generously sharing data on conserved TAD boundaries. Dennis Kostka and Noah Simons provided valuable insight on RNAseq analysis. We also thank members of the Capra Lab who gave helpful feedback throughout this project. We are grateful for the support and resources from the University of Utah Center for High Performance Computing and the Wynton High Performance Compute Cluster at University of California San Francisco. JAC and CMB were funded by National Institutes of Health grant R35GM127087. THW was funded by National Science Foundation grant BCS 1945782.

## 4.15 Author Contributions

Conceptualization, CMB and JAC; Formal Analysis, CMB, SK, ENG, and THW; Writing – Original Draft, CMB and JAC; Writing – Review & Editing, CMB, SK, ENG, EM, KSP, THW, and JAC.

## 5 Supplementary Information

### 5.1 Supplementary Text

#### 5.1.1 *In silico* mutagenesis of chimpanzee lineage-specific variants

We identified lineage-specific 3D-modifying variants in chimpanzee subspecies using *in silico* mutagenesis. We found 78 unique variants with 150 variant-window pairs in central chimpanzees, 337 unique variants with 610 variant-window pairs in eastern chimpanzees, 34,474 unique variants with 64,657 variant-window pairs in Nigeria-Cameroon chimpanzees, and 11,993 unique variants with 22,671 variant-window pairs in western chimpanzees. None of the central or eastern chimpanzee-specific variants yielded divergence > 0.001 when inserted into the reference sequence (**File S5**). This threshold yielded six variants in unique windows among Nigeria-Cameroon chimpanzees; however, the effects were quite small ranging from 0.001–0.008 (**File S5**). Four unique variants resulted in divergence > 0.001 in western chimpanzee, including two that had an effect in both overlapping windows (**File S5**). Of these six windows represented by these variants, only one was previously identified as a western chimpanzee divergent window (**Table S3**). Divergence ranged from 0.001 to 0.009 for all but one of the variants. A C > T mutation (chr2A: 55,039,344) generated a divergence of 0.079 for window chr2A: 54,525,953–55,574,529.

### 5.2 Supplementary Files

**File S1.** This file contains information on the sex, lineage, the number of biallelic SNVs that passed quality filters, and inclusion/exclusion in downstream analyses per individual.

**File S2.** This file contains the results of hierarchical clustering based on 3D divergence per window to identify and assign topologies.

**File S3.** This file contains the outputs from the phenotype enrichment analyses. Results for any trait from the four considered ontologies with at least one gene annotation represented by the 431 genes among the 89 bonobo-chimpanzee divergent windows are included here.

**File S4.** This file contains information on 3D-modifying variants identified from the *in silico* mutagenesis of bonobo-specific variants.

**File S5.** This file contains information on 3D-modifying variants identified from the *in silico* mutagenesis of chimpanzee subspecies-specific variants.

### Supplemental Figures

**Figure S1:**
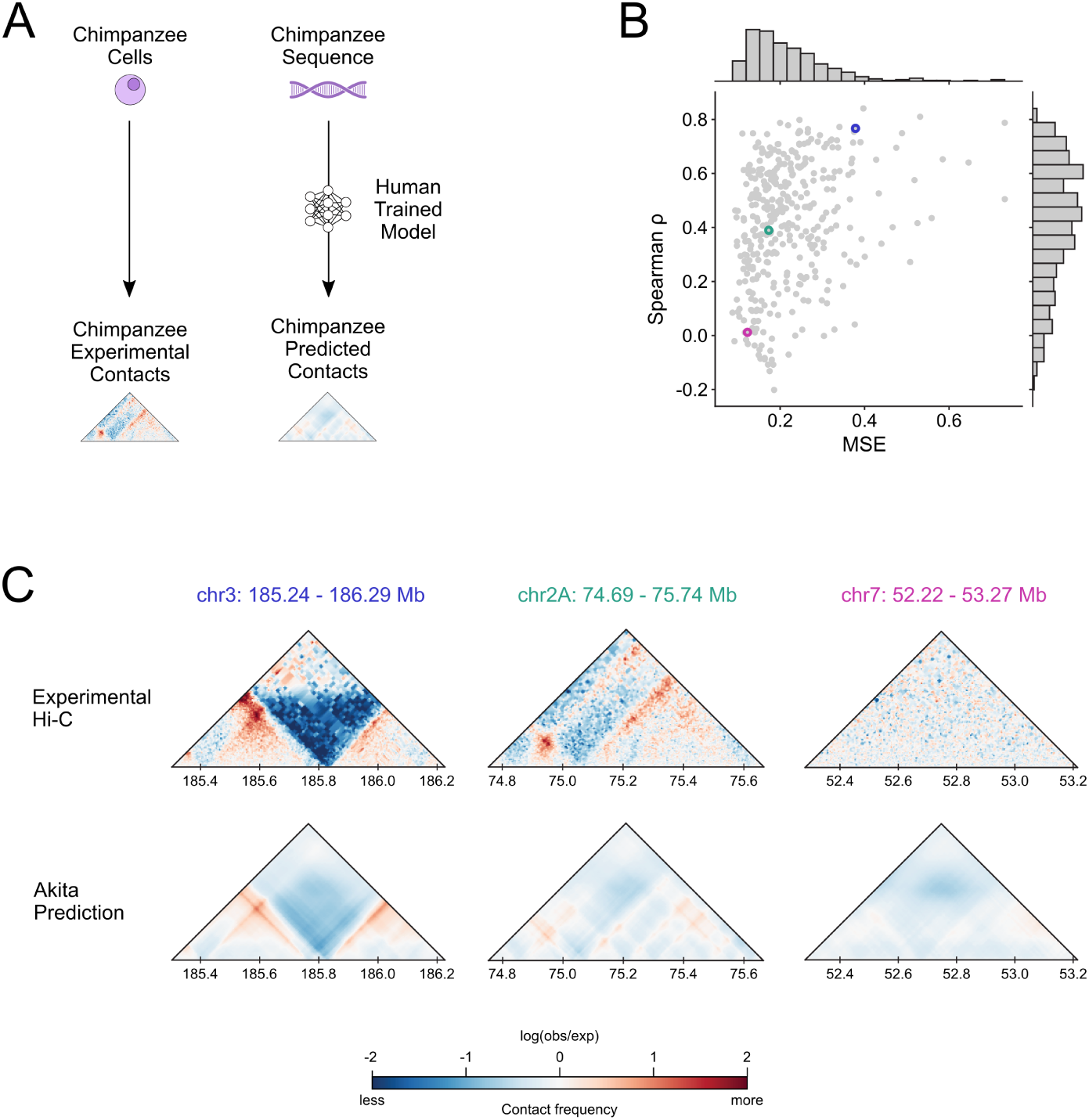
Akita recapitulates the genome folding of chimpanzee in neural progenitor cells. **(A)** Schematic of comparing experimental chromatin contacts to predicted chromatin contacts. Hi-C data were from chimpanzee neural progenitor cells. Predictions were acquired from the HFF output of the human-trained Akita model on panTro6 sequence. We compared chimpanzee regions (N = 368) lifted over from the human held-out test set in Fudenberg et al., 2020. **(B)** The mean squared error (MSE) versus Spearman *ρ* between the experimental Hi-C contact map and Akita prediction for each of the test set windows. **(C)** Experimental and predicted contact maps for three example regions highlighted in **B** with blue, green, and pink circles.

**Figure S2:**
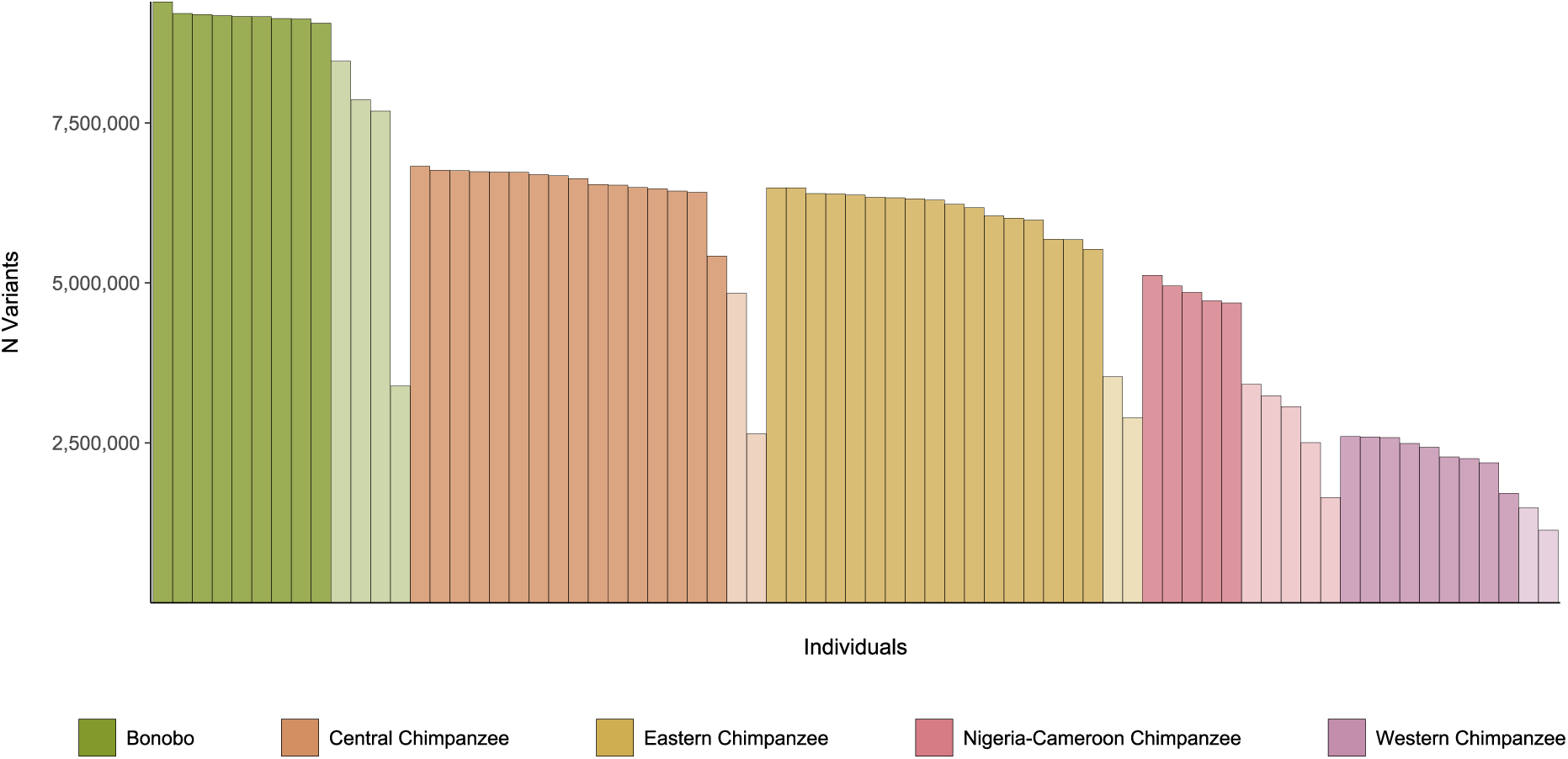
Variants per individual. The number of SNVs inserted to the panTro6 reference sequence per individual. Color indicates the lineage per individual. Individuals excluded from final analysis (N = 15) are shaded lighter than included individuals. See **File S1** for details.

**Figure S3:**
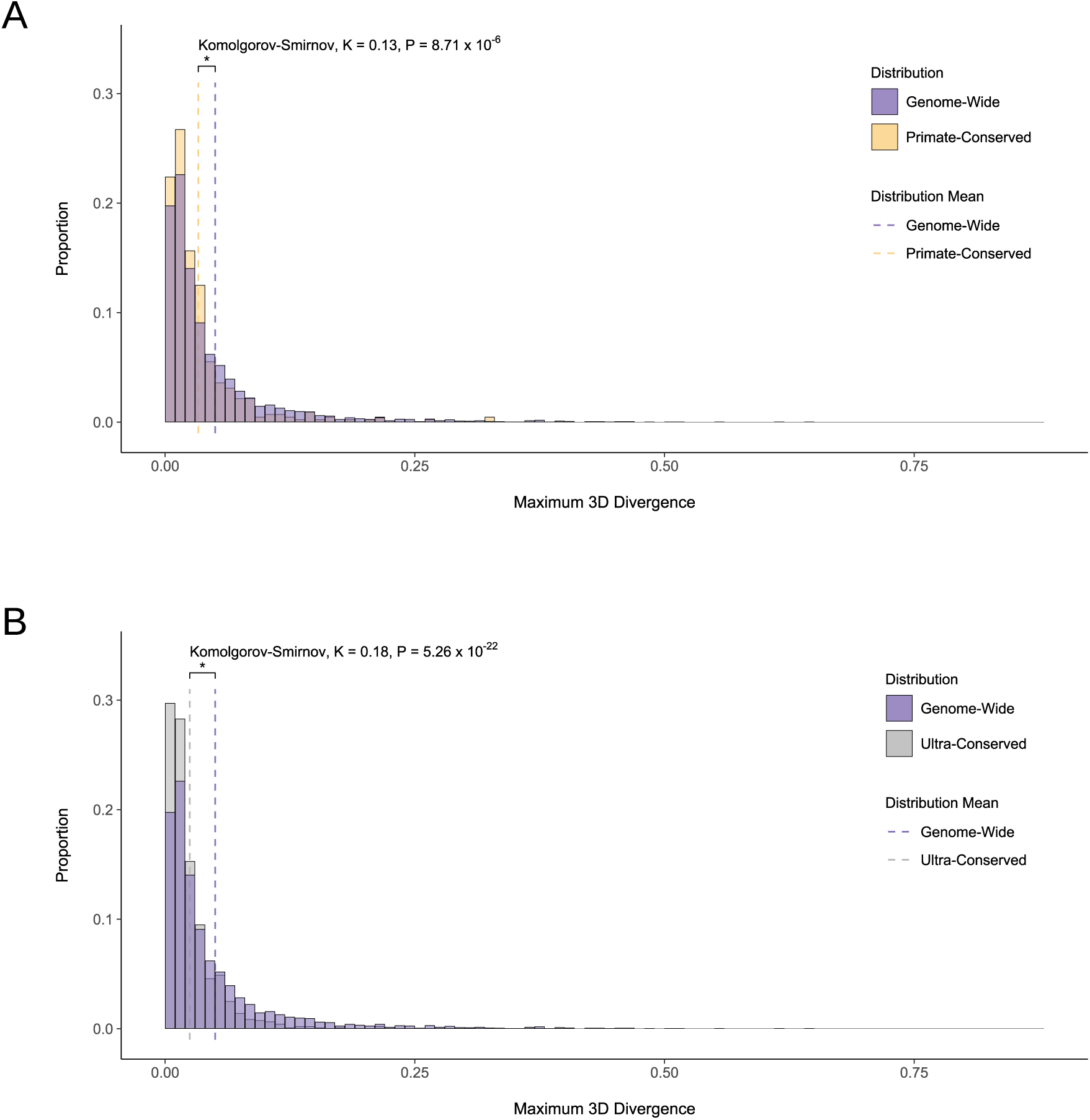
Experimentally-validated conserved regions of the 3D genome are minimally 3D divergent among bonobos and chimpanzees. **(A)** The distribution of maximum 3D divergence for all windows (N = 4,420) and the most central window intersecting primate-conserved TAD boundaries from Okhovat et al., 2023 (N = 415). Distributions are shown in 0.01 divergence bins and the dashed line indicates the distribution means. The genome-wide distribution and mean are shown in purple and the primate-conserved distribution and mean shown in yellow. **(B)** The distribution of maximum 3D divergence for all windows (N = 4,420) and the most central window intersecting ultra-conserved TAD boundaries from Okhovat et al., 2023 (N = 915). Distributions are shown in 0.01 divergence bins and the dashed line indicates the distribution means. The genome-wide distribution and mean are shown in purple and the ultra-conserved distribution and mean are shown in gray.

**Figure S4:**
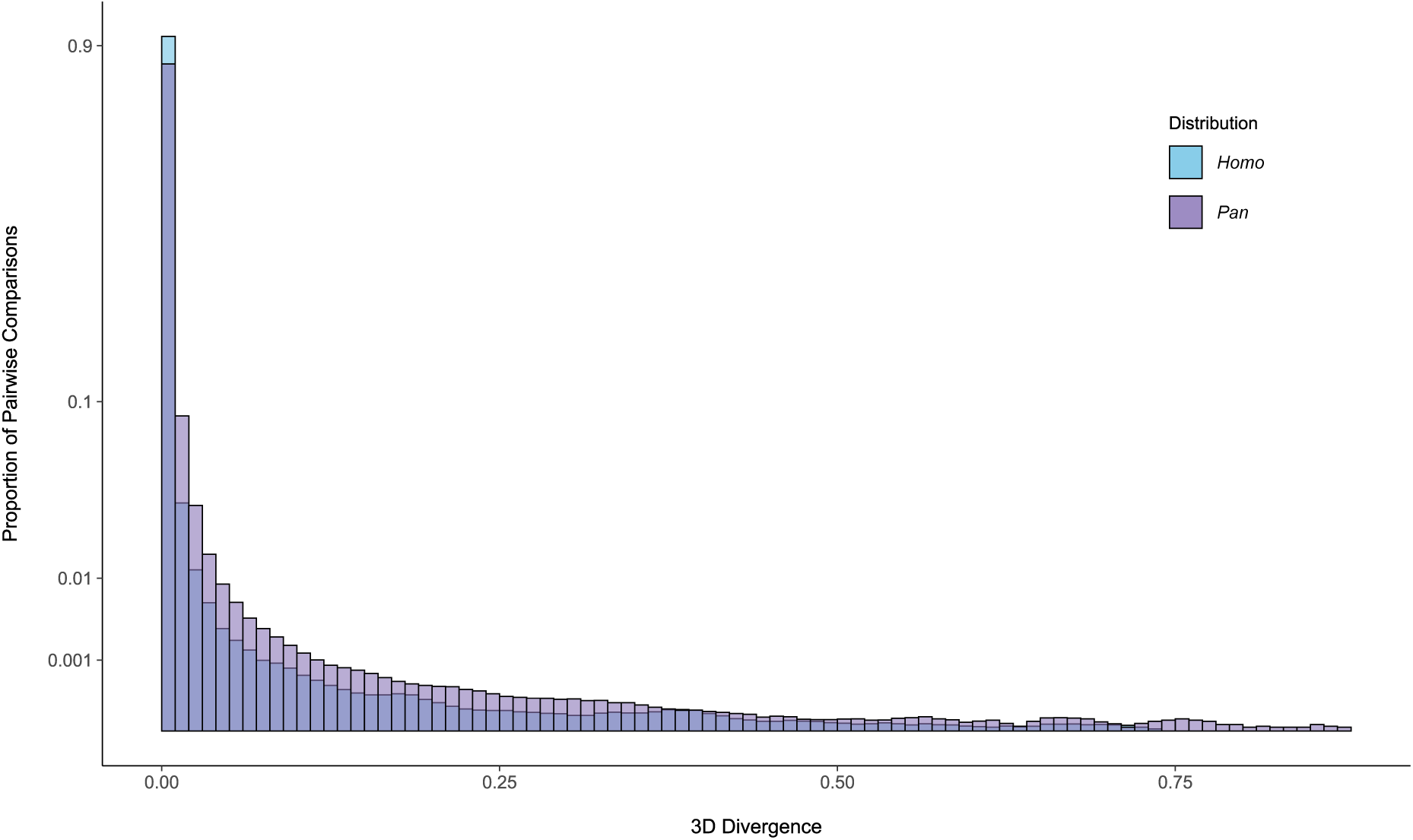
3D divergence within *Pan* is greater than within genetically diverse modern humans. The distribution of 3D divergence among autosomes in 0.01 divergence bins in 56 bonobos and chimpanzees from the present study (purple) compared to 130 modern humans from Thousand Genomes Project (1KG) (Gilbertson et al., in prep) (blue). The *Pan* distribution comprises 6,574,260 comparisons and the modern human distribution comprises 40,860,105 comparisons. The modern human sample consists of five individuals from each of the 26 1KG subpopulations. *Pan* 3D divergence is significantly higher (mean = 0.008) than the modern human distribution (mean = 0.003) (Komolgorov-Smirnov, K = 0.329, P = 2.23 *×* 10^-308^). Note the y-axis is cube root transformed.

**Figure S5:**
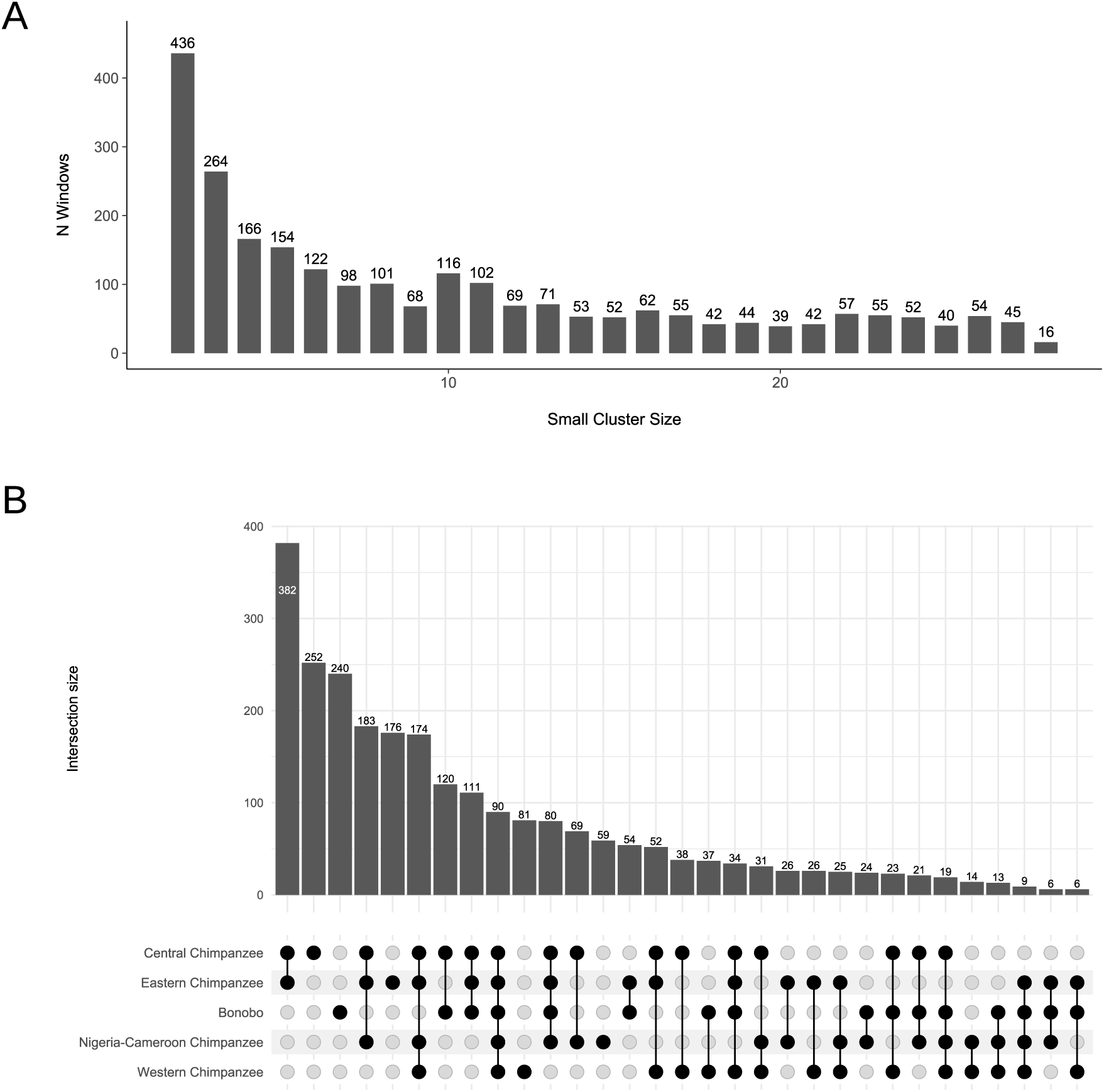
Windows with a multiple divergent individuals window topology commonly feature a small cluster featuring few individuals representing the three most genetically diverse *Pan* lineages. **(A)** The small cluster size distribution among windows with a multiple divergent individuals topology. **(B)** The distribution of lineages represented in each small cluster. Bars indicate the number of windows with a given small cluster size and the dot matrix indicates the lineages represented.

**Figure S6:**
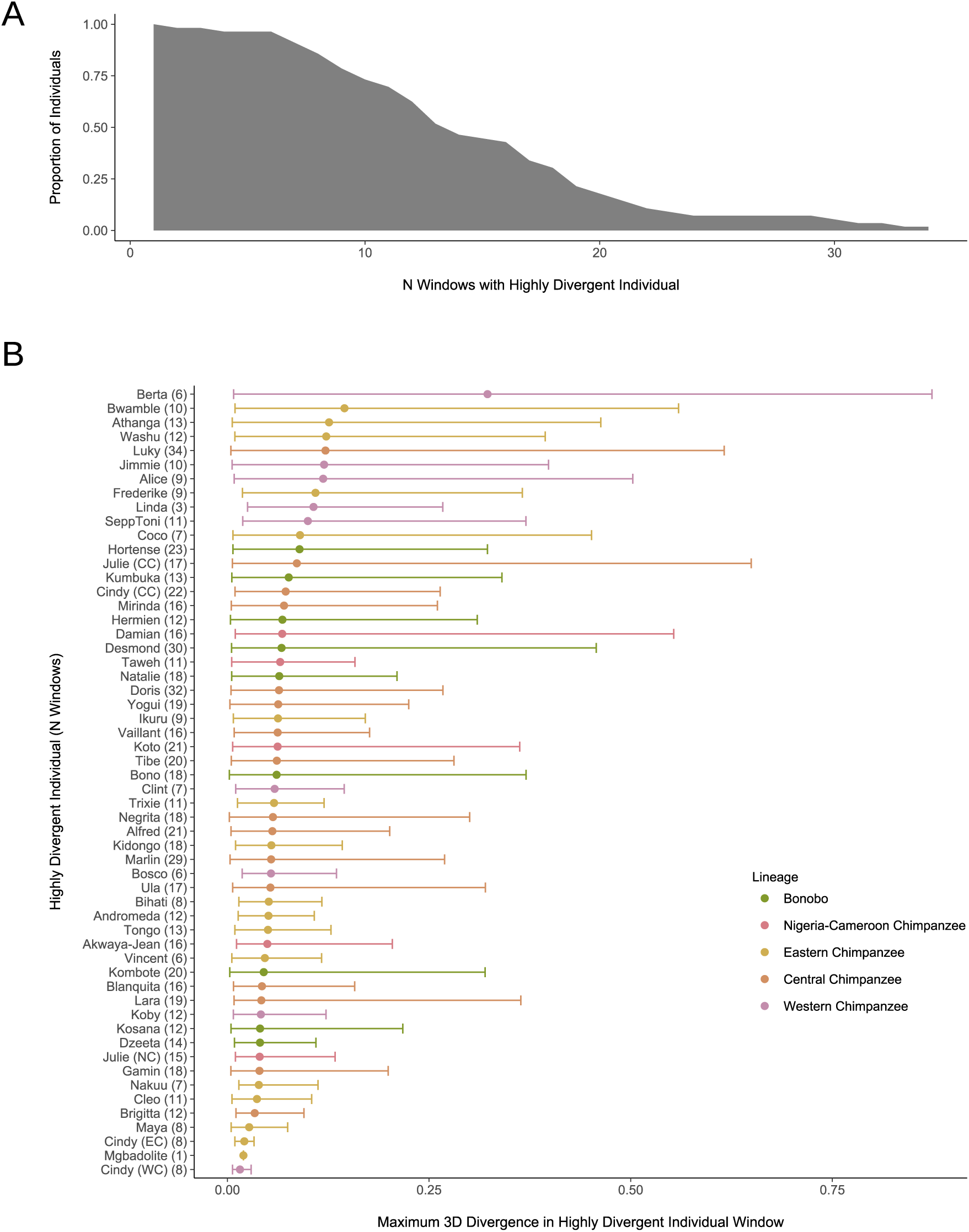
Highly divergent individual windows are common across individuals. **(A)** The inverse cumulative density of individuals with *≥* N windows where they are the divergent individual. **(B)** The distribution of 3D divergence maxima for each individual’s set of highly divergent windows. The minimum, mean, and maximum are indicated by lower error bar, point, and upper error bar, respectively. Individuals are ordered by decreasing mean. While no lineages contain individuals with same name, two names are present among multiple lineages: “Cindy” and “Julie”. These individuals are distinguished by lineage using parentheses: CC = central chimpanzee, EC = eastern chimpanzee, NC = Nigeria-Cameroon chimpanzee, and WC = western chimpanzee. The number of windows per individual where they are divergent is displayed in parentheses.

**Figure S7:**
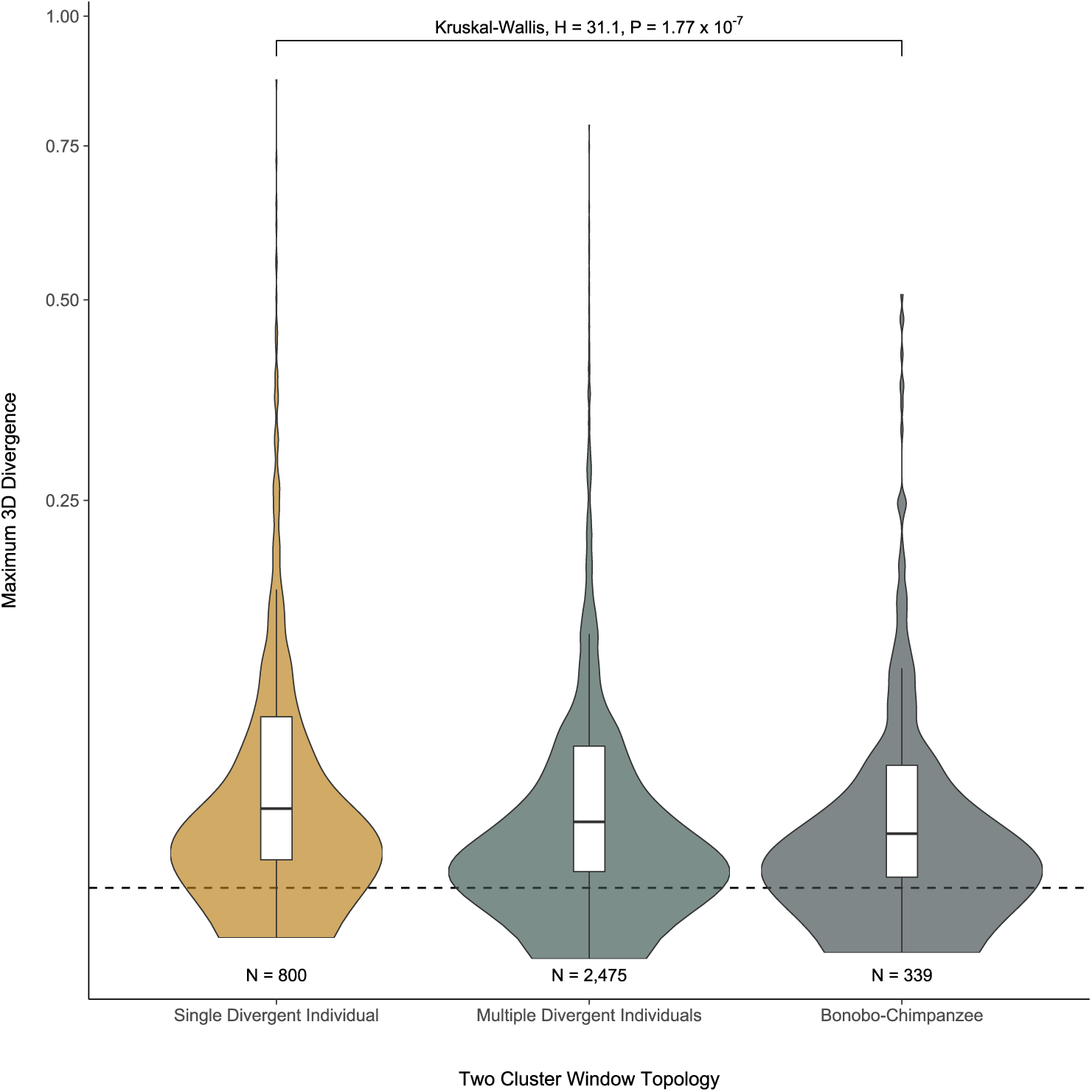
Two cluster topologies significantly differ in maximum 3D divergence. The distribution of maximum 3D divergence per window stratified by two cluster window topologies: highly divergent individual, multiple divergent individuals, and bonobo-chimpanzee clustering windows. Highly divergent individual clustering windows are more 3D divergent (mean = 0.067) than multiple divergent individuals (mean = 0.053) or bonobo-chimpanzee (mean = 0.049) clustering windows. Violin plots show density and the boxplots display the median and IQR, with the upper whiskers extending to the largest value *≤* 1.5 x IQR from the 75th percentile and the lower whiskers extending to the smallest values *≤* 1.5 x IQR from the 25th percentile. Outliers are not displayed in the boxplots. The horizontal dashed line indicates 3D divergence of 0.01. Note the y-axis is square root transformed.

**Figure S8:**
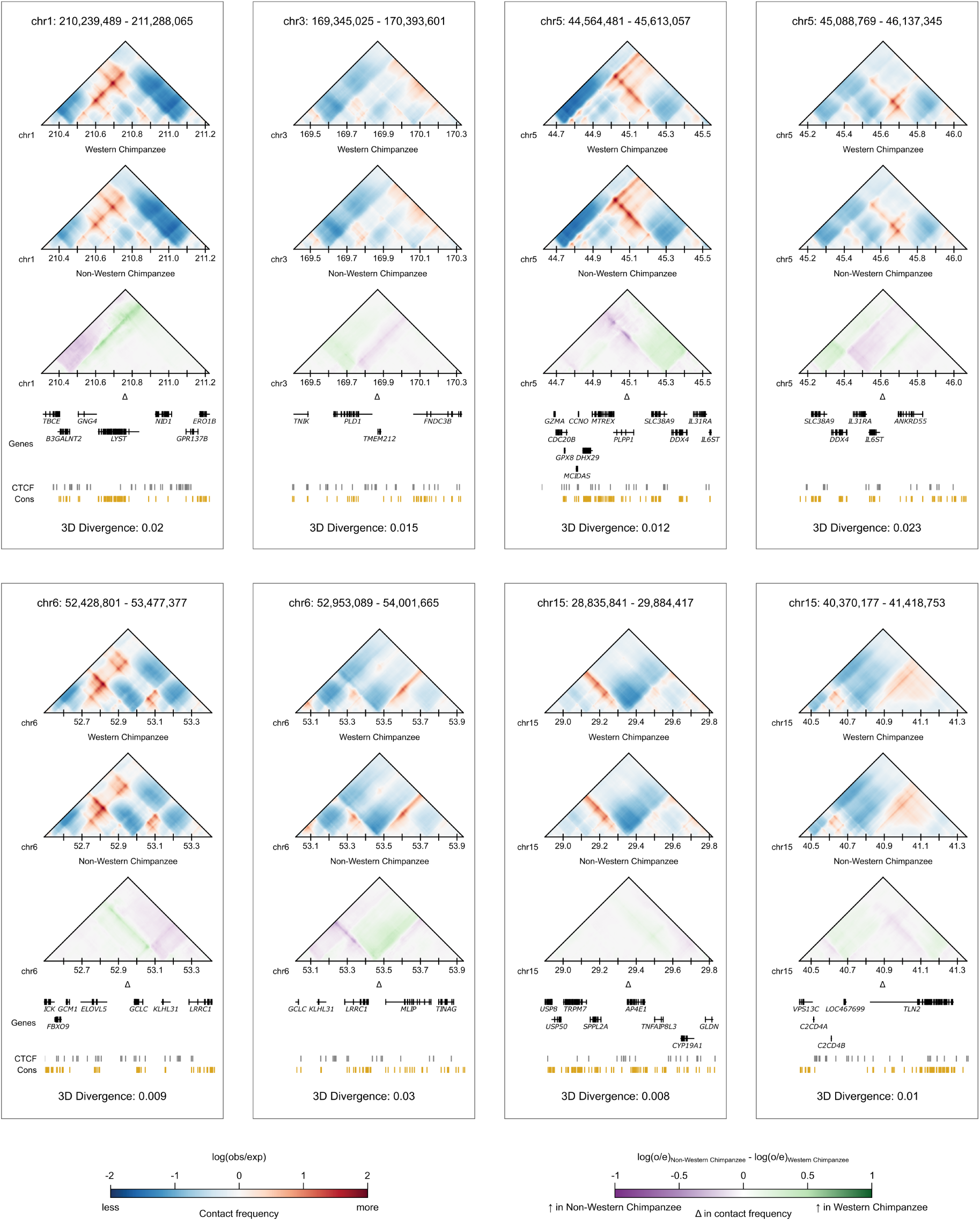
Western chimpanzees cluster separately to all other bonobos and chimpanzees at eight genomic windows. Contact maps for a western chimpanzee, a non-western chimpanzee, and the contact difference (Δ) at each of the eight windows where western chimpanzees clustered separately to all other bonobos and chimpanzees. The most divergent pair is shown per window. Maps are annotated with genes, CTCF peaks from Schwalie et al., 2013 and phastCons conserved elements from the UCSC Genome Browser. The individual western chimpanzees shown in these maps are SeppToni, Jimmie, Koby, Jimmie, Bosco, Bosco, Bosco, and Clint (L to R, top to bottom). The individual non-western chimpanzees shown in these maps are Gamin, Tongo, Hermien, Cindy (EC), Julie (CC), Kumbuka, Mirinda, and Andromeda (L to R, top to bottom).

**Figure S9:**
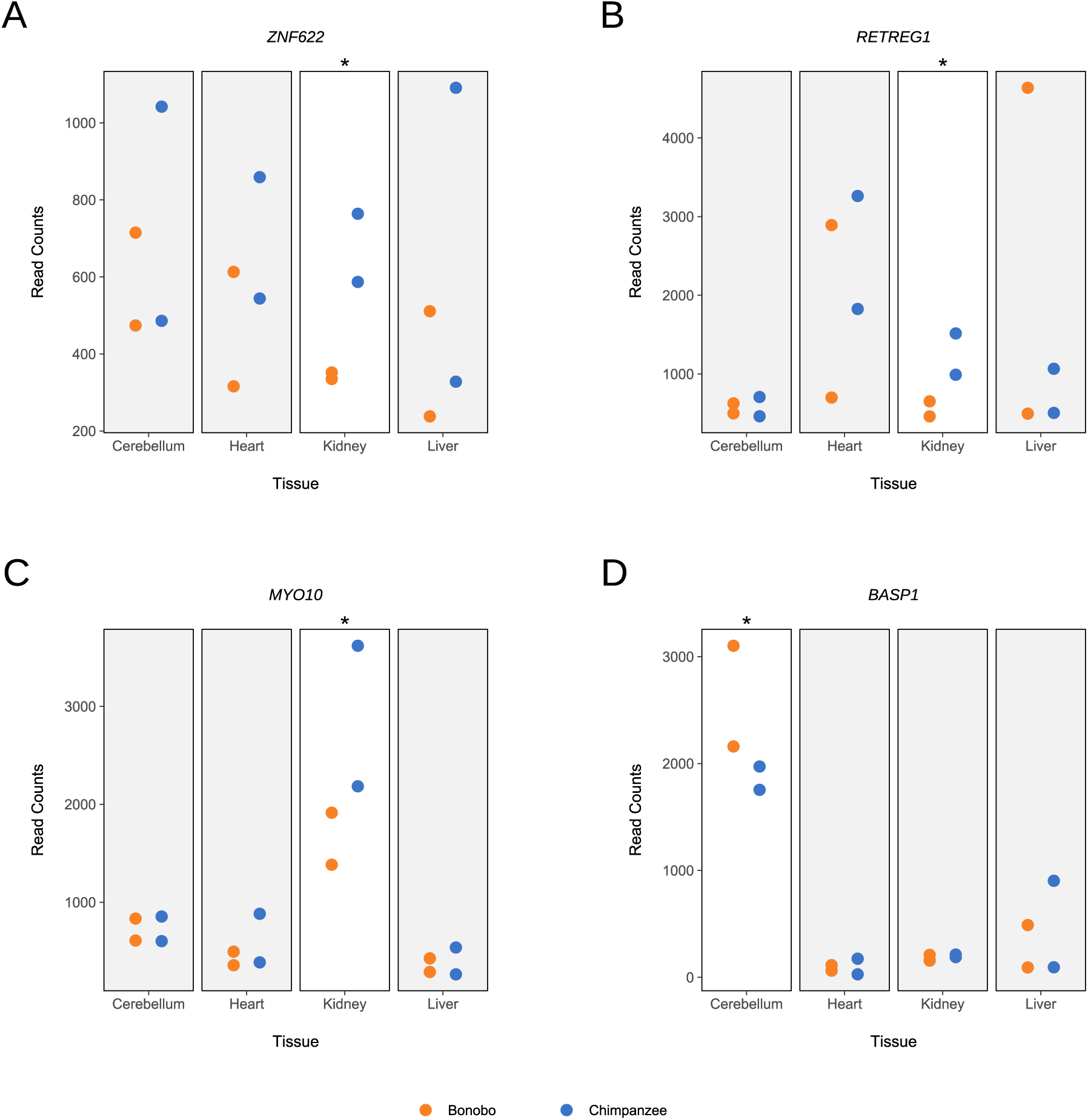
Bonobos and chimpanzees exhibit tissue-specific expression differences at a bonobo-chimpanzee divergent window. mRNA read counts for all genes intersecting the chr5: 16,252,929 - 17,301-505 bonobo-chimpanzee divergent window: **(A)** *ZNF622*, **(B)** *RETREG1*, **(C)** *MYO10*, and **(D)** *BASP1*. We display the four tissues with two samples per species from Brawand et al., 2011, indicating species by color. Genes are ordered by increasing start coordinate. Tissues with a species difference in expression are not shaded and noted by an asterisk. See **Methods** for details on RNAseq processing and quantifying reads.

**Figure S10:**
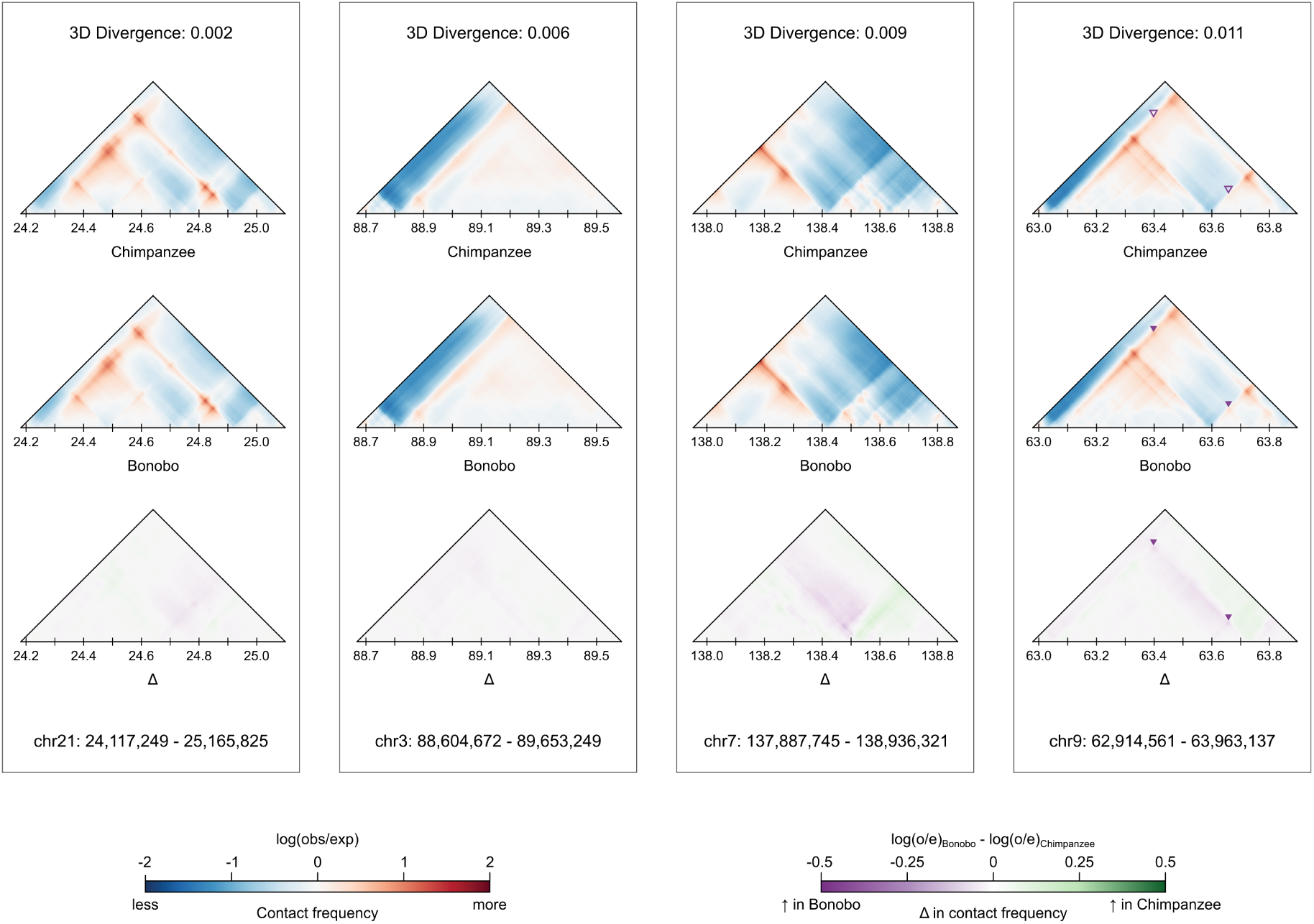
Contact map differences between bonobos and chimpanzees are more subtle among windows with an interspecific topology and low 3D divergence minima. Contact maps for a chimpanzee, a bonobo, and the contact difference (Δ) for the interspecific comparison with the lowest 3D divergence at four different windows characterized by an interspecific topology. The individual bonobos shown in these maps are Bono, Kosana, Hermien, and Dzeeta (L to R). The individual chimpanzees shown in these comparisons are Koto, Kidongo, Linda, and Luky (L to R).

**Figure S11:**
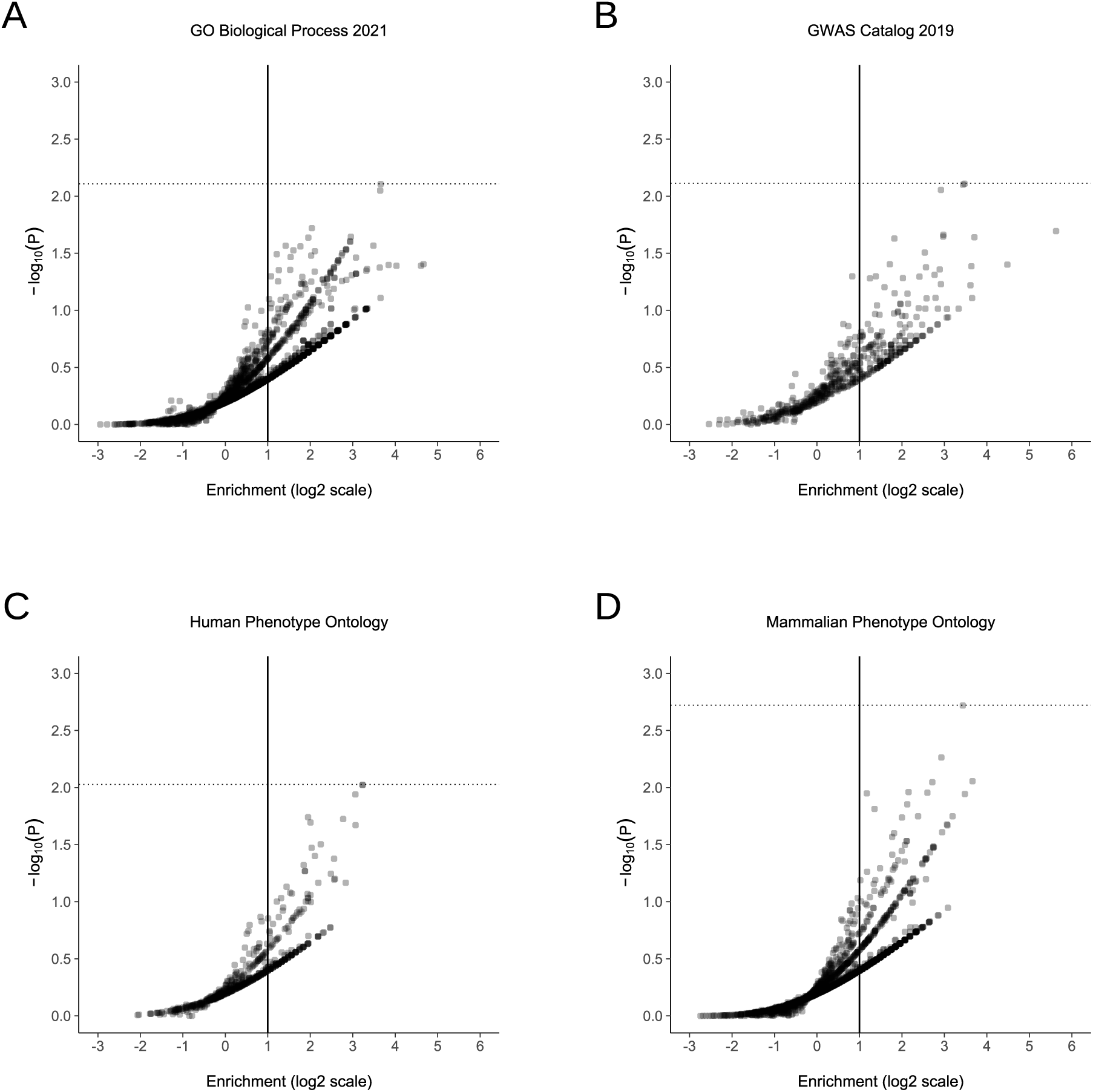
Bonobo-chimpanzee divergent windows are not enriched for genes underlying biological processes, human disease, or mammalian phenotypes. **(A)** Enrichment of genes associated with 2,135 phenotypes in the GO Biological Process 2021 catalog among windows with a bonobo-chimpanzee topology. Each point represents a phenotype. Enrichment and p-values were calculated from a one-sided permutation test based on an empirical null distribution generated from 10,000 shuffles of maximum Δ across the entire dataset (Methods). The vertical solid line indicates no enrichment and the horizontal dotted line represents the false-discovery rate (FDR) corrected p-value threshold at FDR = 0.05. See **File S3** for all phenotype enrichment results. **(B)** Enrichment of genes associated with 552 phenotypes in the GWAS Catalog 2019 catalog among windows with a bonobo-chimpanzee topology. Data were generated and visualized as in **A**. **(C)** Enrichment of genes associated with 621 phenotypes in the Human Phenotype Ontology among windows with a bonobo-chimpanzee topology. Data were generated and visualized as in **A**. **(D)** Enrichment of genes associated with 1,740 phenotypes in the Mammalian Phenotype Ontology among windows with a bonobo-chimpanzee topology. Data were generated and visualized as in **A**.

**Figure S12:**
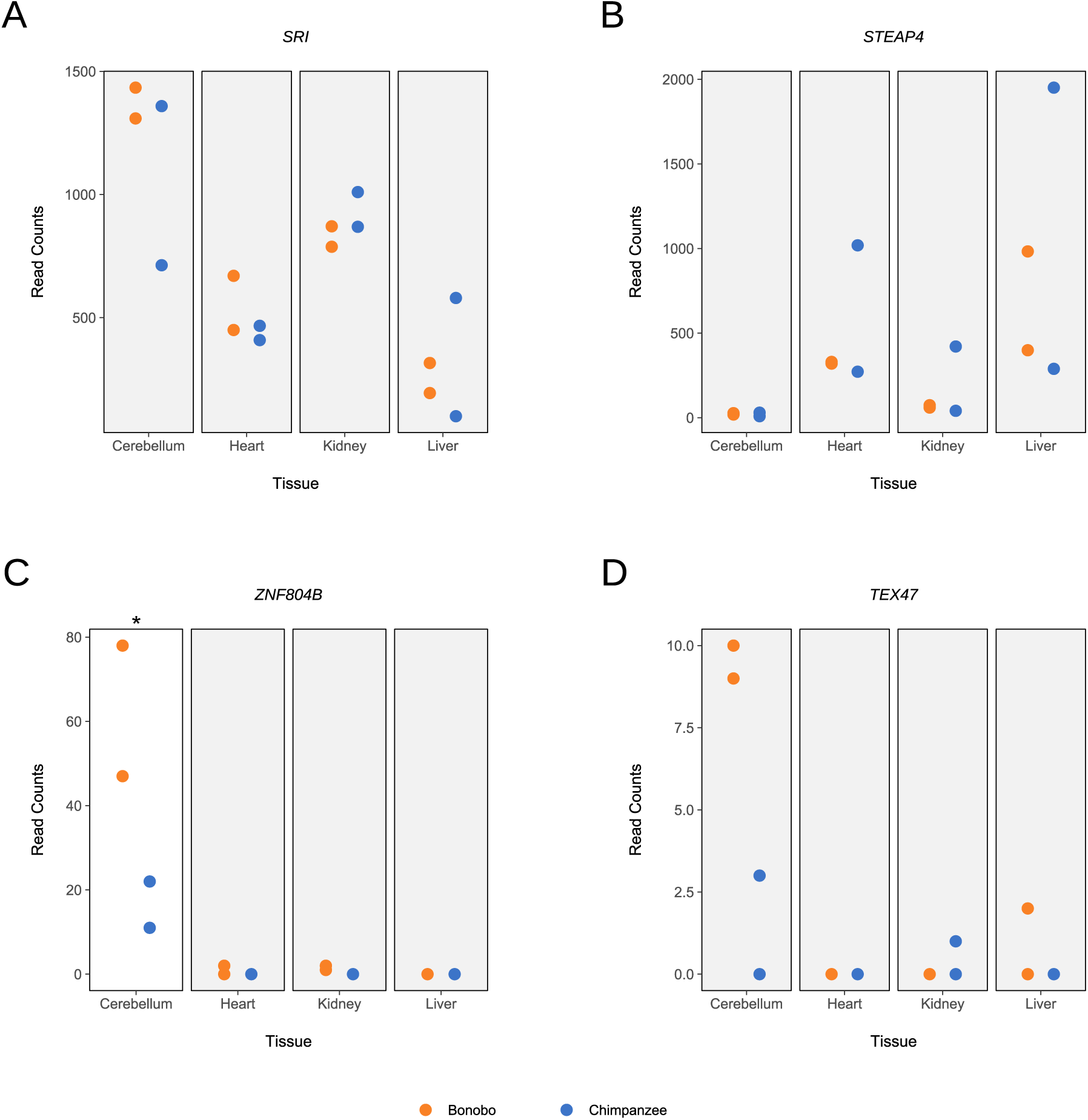
Bonobos and chimpanzees exhibit tissue-specific expression differences at a bonobo-chimpanzee divergent window. mRNA read counts for genes near the species difference in genome folding within the chr7: 83,886,081 - 84,934,657 bonobo-chimpanzee divergent window: **(A)** *SRI*, **(B)** *STEAP4*, **(C)** *ZNF804B*, and **(D)** *TEX47*. We display the four tissues with two samples per species from Brawand et al., 2011, indicating species by color. Genes are ordered by increasing start coordinate. Tissues with a species difference in expression are not shaded and noted by an asterisk. Any tissue with a read count of zero for one or more samples were not considered and are shaded. See **Methods** for details on RNAseq processing and quantifying reads.

**Figure S13:**
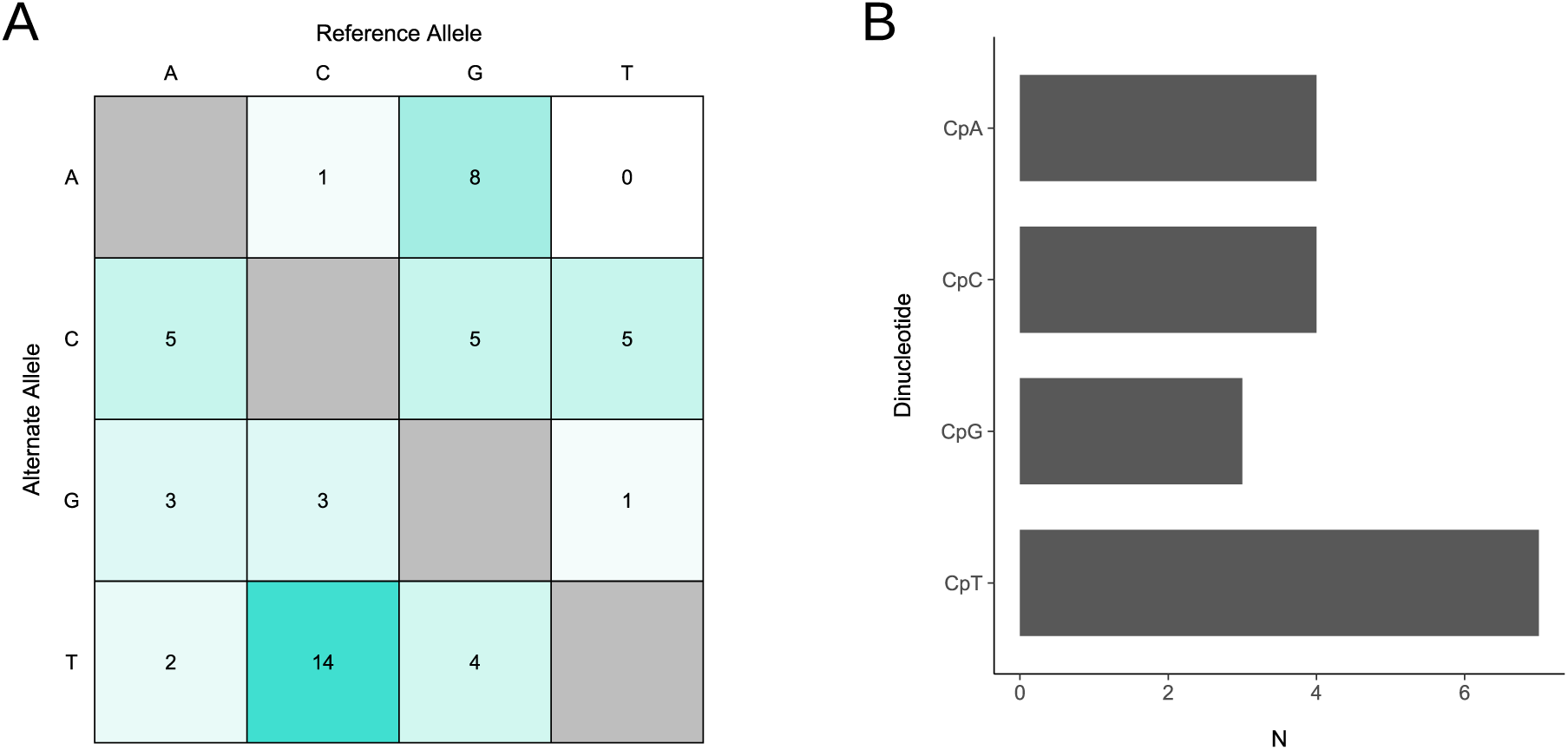
3D-modifying variant mutations are non-randomly distributed and not driven by GC-biased gene conversion. **(A)** The mutation matrix for 51 3D-modifying variants. Cells are shaded by frequency. **(B)** Dinucleotide context counts for the 18 mutations with a reference “C” allele.

**Figure S14:**
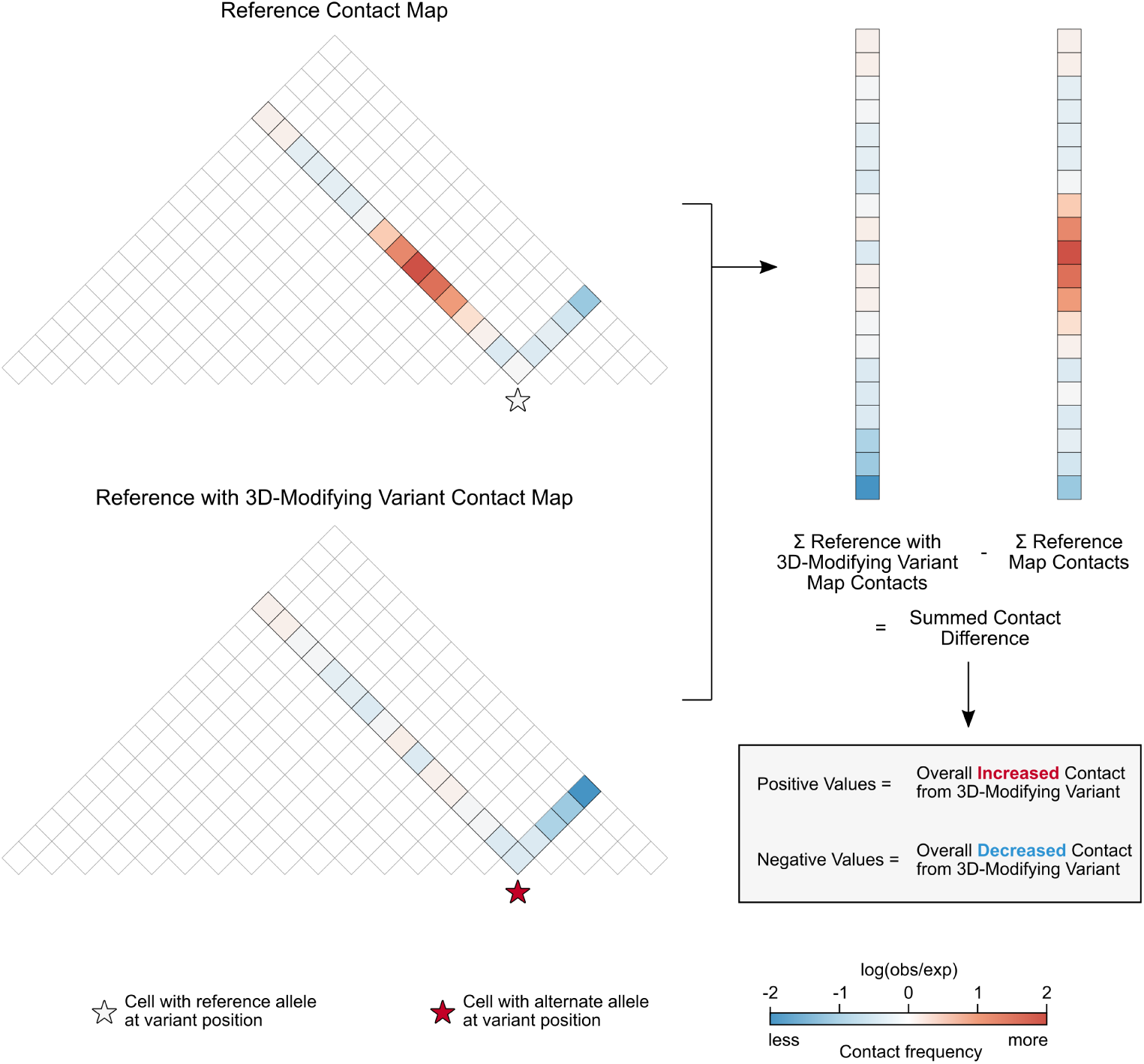
Summing contact differences captures the overall effect of 3D-modifying variants. To quantify the effect of each 3D-modifying variant, we calculated the summed contact difference for the reference map and the reference with 3D-modifying variant map. We added the contact frequencies for all 2,048 bp cells that represented pairwise contact between the cell containing the variant and all other cells in the window (colored diagonals). The example here yields a summed contact frequency for 20 cells, while sums of the empirical data are calculated from 448 cells. Frequencies are illustrated using color and cells with pairwise contacts that do not involve the variant cell are not colored. We subtracted the summed contacts of the reference map from the summed contacts of the reference with variant map. Thus, a positive summed contact difference indicates overall increased contact from the 3D-modifying variant, whereas negative values indicates overall decreased contact.

**Figure S15:**
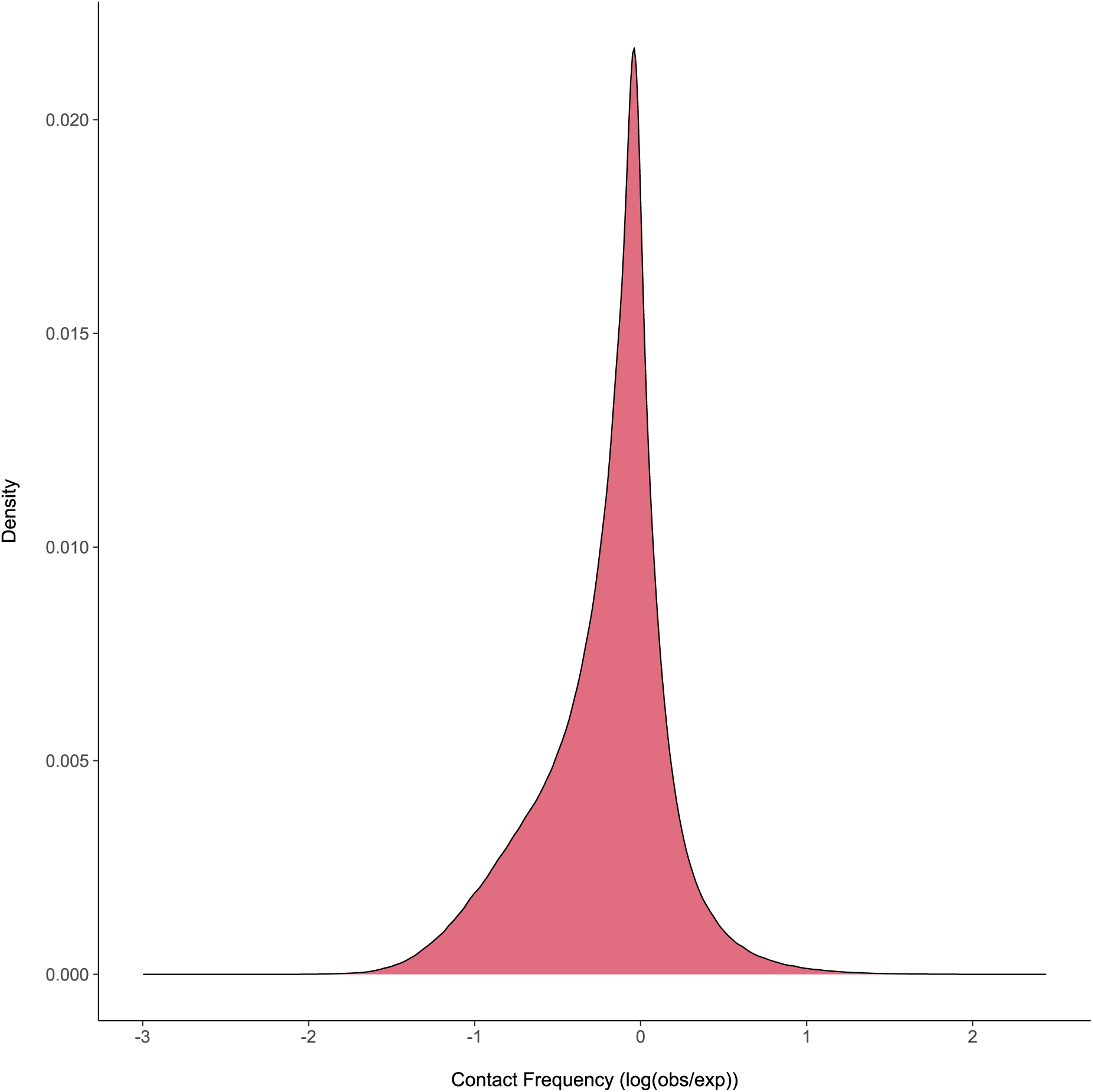
Most predicted contact frequencies fall between -2 and 2. A distribution of predicted contact frequencies sampled from 2,800,000 frequencies—50,000 randomly chosen values per individual included in the final analysis (N = 56).

### Supplemental Tables

**Table S1:** Median 3D divergence is positively associated with effective population size for within lineage comparisons. We stratified comparisons by the *Pan* lineages represented in each pair and computed the median 3D divergence for comparisons made within the same lineage. Effective population size or *N_e_* listed here is the median value from Prado-Martinez et al., 2013.

| Lineage | $N_e$ | Median 3D Divergence |
| --- | --- | --- |
| Central Chimpanzee | 36,550 | 0.0023 |
| Eastern Chimpanzee | 29,600 | 0.0016 |
| Nigeria-Cameroon Chimpanzee | 27,750 | 0.0013 |
| Bonobo | 17,850 | 0.0008 |
| Western Chimpanzee | 14,650 | 0.0006 |

**Table S2:**
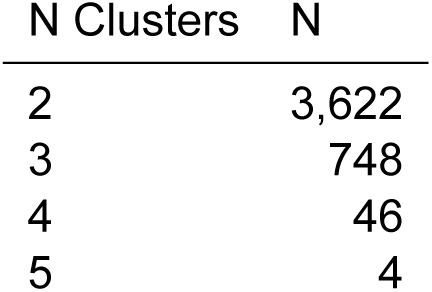
3D divergence in most genomic windows resulted in two clusters after hierarchical clustering. The number of genomic windows with a given number of clusters.

**Table S3:** Western chimpanzees cluster separately to all other bonobos and chimpanzees at eight genomic windows. The chromosome and position start (1-based), maximum 3D divergence, and overlapping genes for eight windows where western chimpanzees cluster separately to all other bonobos and chimpanzees.

| Chr | Window Start | Maximum Divergence | Genes |
| --- | --- | --- | --- |
| chr1 | 210,239,489 | 0.020 | ARID4B, B3GALNT2, ERO1B, GGPS1, GNG4, GPR137B, LYST, NID1, TBCE |
| chr3 | 169,345,025 | 0.015 | FNDC3B, PLD1, TMEM212, TNIK |
| chr5 | 44,564,481 | 0.012 | CCNO, CDC20B, DDX4, DHX29, ESM1, GPX8, GZMA, GZMK, IL31RA, IL6ST, MCIDAS, MTREX, PLPP1, SLC38A9 |
| chr5 | 45,088,769 | 0.023 | ANKRD55, DDX4, IL31RA, IL6ST, PLPP1, SLC38A9 |
| chr6 | 52,428,801 | 0.009 | ELOVL5, FBXO9, GCLC, GCM1, GSTA4, ICK, KLHL31, LRRC1 |
| chr6 | 52,953,089 | 0.030 | GCLC, KLHL31, LRRC1, MLIP, TINAG |
| chr15 | 28,835,841 | 0.008 | AP4E1, CYP19A1, DMXL2, GLDN, SPPL2A, TNFAIP8L3, TRPM7, USP50, USP8 |
| chr15 | 40,370,177 | 0.010 | C2CD4A, C2CD4B, LOC467699, TLN2, VPS13C |

